# Trained Immunity Causes Myeloid Cell Hypercoagulability

**DOI:** 10.1101/2024.07.16.603679

**Authors:** Aisling M. Rehill, Seán McCluskey, Anna E. Ledwith, Tristram A.J. Ryan, Gemma Leon, Hugo Charles-Messance, Edmund H. Gilbert, Emily A. Day, Corrina McMahon, James S. O’Donnell, Annie M. Curtis, Luke A.J. O’Neill, Frederick J. Sheedy, Roger J.S. Preston

## Abstract

Venous thromboembolism is common in individuals with chronic inflammatory diseases, but the pathogenic basis for this increased thrombotic risk remains poorly understood. Myeloid cell ‘trained immunity’ describes persistent innate immune cell memory arising from prior exposure to an inflammatory stimulus, leading to an enhanced immune response to subsequent unrelated stimuli. We identify enhanced myeloid cell prothrombotic activity as a novel maladaptive consequence of trained immunity. LPS stimulation of murine bone marrow-derived macrophages trained previously with either β-glucan or free haem exhibited significantly enhanced procoagulant and antifibrinolytic gene expression and activity compared to macrophages stimulated with LPS alone. The β-glucan training-mediated increase in activated myeloid cell procoagulant activity was mediated by enhanced acid sphingomyelinase-mediated tissue factor (TF) functional decryption. Furthermore, pre-treatment with methyltransferase and acetyltransferase inhibitors to erase epigenetic marks associated with innate immune memory diminished trained macrophage TF gene expression in β-glucan-trained macrophages. Functional analysis of splenic monocytes isolated from β-glucan-trained mice revealed enhanced procoagulant activity up to 4 weeks after β-glucan administration compared to monocytes from control mice over the same time period. Remarkably, monocyte procoagulant activity increased proportionately with time since β-glucan administration, before plateauing at 4 weeks. Furthermore, haematopoietic progenitor cells and bone marrow interstitial fluid isolated from β-glucan-trained mice possessed enhanced procoagulant activity compared to control mice. Trained immunity and associated metabolic perturbations may therefore represent novel therapeutic vulnerabilities in immunothrombotic disease development, opening new avenues for targeted intervention.

## INTRODUCTION

The risk of venous thromboembolism is significantly elevated in patients with inflammatory disorders and haematological malignancies, which are often characterised by extended periods of chronic inflammation. These conditions, which include sickle cell disease^1^, inflammatory bowel disease^2^ and myeloproliferative disorders^3^, amongst others, have a high worldwide prevalence and significant individual burden. Despite the elevated risk of thrombosis in affected individuals, the mechanistic basis for how chronic inflammatory disease might also promote haemostatic dysregulation in the long term is not understood. Elucidating the pathophysiological mechanisms that underlie increased thrombotic risk in individuals with inflammatory diseases is crucial to the development of new treatment options and prevention strategies.

Aberrant activation of peripheral and tissue-resident innate immune cells is strongly associated with the development of immunothrombosis^4,5^. In addition to their acute response to pathogens, innate immune cells can retain a ‘memory’ of past exposure to pathogens or inflammatory stimuli, predisposing these cells to a heightened pro-inflammatory response upon re-exposure to a non-specific inflammatory event. This process is termed ‘*trained immunity*’ or ‘*innate immune memory’*^6^. Unlike an adaptive immune response, this phenomenon is not antigen-specific, and has likely evolved to enable innate immune cells to mount a robust response to subsequent pathogenic exposure. Well-characterised mediators of trained immunity include the Bacillus Calmette-Guerin (BCG) vaccine^7^, *Candida albicans*^8^ and β-glucan^9^, a polysaccharide cell wall component. Moreover, sterile inflammation, caused by oxidised low-density lipoprotein (LDL)^10^ or free haem^11^, can also confer innate immune memory.

To imprint innate immune memory, myeloid cells undergo extensive functional reprogramming via epigenetic and metabolic changes^12^. β-glucan-trained macrophages exhibit enhanced H3 histones trimethylated at lysine 4 (H3K4me3) and H3 histones acetylated at lysine 27 (H3K27ac) at the promotor and/or enhancer regions of key pro-inflammatory and metabolic genes^9,13,14^. Profound changes in myeloid cell metabolism, including heightened glycolytic pathway activity, also accompany the phenotypic changes that arise from innate immune memory^14,15^. *In vivo*, innate immune memory is mediated by phenotypic reprogramming of haematopoietic stem and progenitor cells (HSPCs). HSPC populations in the bone marrow (BM) rapidly proliferate in response to systemic inflammation or severe infection response^8,9^. The induction of trained immunity promotes the expansion of HSPCs with a bias towards myelopoiesis, and the myeloid cells generated have a lowered threshold for pro-inflammatory activity upon re-stimulation^16,17^.

Although trained immunity is likely beneficial in boosting innate immune responses by generating heightened pro-inflammatory responses to subsequent pathogen exposure, maladaptive induction of trained immunity has also been linked to an increased propensity to develop and exacerbate chronic inflammatory disease. For example, metabolic dysfunction arising from a high-fat diet leads to HSPC myelopoietic bias and heightened myeloid cell pro-inflammatory activity in mice^18^. Notably, maladaptive BM-trained immunity induced by periodontitis in mice exacerbates subsequent disease severity and pro-inflammatory activity when the same mice experience inflammatory arthritis, and vice-versa^19^. Interestingly, these data indicate unrelated inflammatory disease activity can predispose to, or exacerbate, the development and severity of an unrelated inflammatory disease via long-term HSPC reprogramming and generation of myeloid cells with enhanced pro-inflammatory preparedness.

We still know little about how chronic or recurring inflammation shapes thrombotic risk in the long term. We hypothesised that this phenomenon may arise from individual inflammatory events that reprogram HSPC generation, leading to the synthesis of myeloid cells with a lowered threshold for induction of immunothrombotic activity. We demonstrate that exposure of monocytes and macrophages to mediators of trained immunity causes epigenetic and metabolic reprogramming, accelerating acid sphingomyelinase (ASMase)-mediated tissue factor decryption and enhancing myeloid cell antifibrinolytic activity. Moreover, we demonstrate that the induction of innate immune memory in mice causes BM reprogramming, which promotes myeloid cell and plasma procoagulant activity for weeks after the original training event.

## RESULTS

### β-glucan-mediated training immunity results in enhanced macrophage procoagulant gene expression

In keeping with previous reports, β-glucan-treated bone marrow-derived macrophages (BMDMs) released significantly more TNFα upon secondary LPS stimulation than when treated with LPS alone and exhibited increased glycolytic activity. **(Supplementary Figure 1a-c)**. To establish whether gene expression of coagulation-associated genes was altered in macrophages by prior induction of innate immune memory, we performed bulk RNA-seq analysis using naïve and LPS-treated BMDMs that were previously exposed to either β-glucan or cell culture media one week prior (**Figure 1a**). DEG analysis of β-glucan-trained BMDMs showed significantly increased expression of the *TNF* and *IL6* genes (encoding TNFα, IL-6 cytokines) than untrained LPS-treated BMDMs, in keeping with previous studies^14,20^ **(Figure 1b, Supplementary Figure 2 and Supplementary Table 2 and 3)**. Interestingly, we noted that β-glucan training also increased the expression of several coagulation-associated genes, including *F3*, *Serpine1* and *Plau,* which encode tissue factor (TF), plasminogen activator inhibitor 1 (PAI-1) and urokinase plasminogen activator (uPA) respectively **(Figure 1b and 1d, Supplementary Table 2**). GO enrichment analysis of LPS-treated BMDMs that were previously treated with β-glucan demonstrated significantly upregulated expression of gene sets associated with ‘coagulation’ and ‘haemostasis’ compared to BMDMs stimulated with LPS alone **(Figure 1c and Supplementary Figure 2b)**. Further qPCR analysis of re-stimulated β-glucan-trained BMDMs demonstrated *F3* **(Figure 1e)** and *Serpine1* **(Figure 1f)** gene expression was significantly increased compared to LPS-treated BMDMs, such that, relative to naïve BMDMs, *F3* expression in trained BMDMs was increased 31-fold, compared to only a 3.5-fold increase in LPS-treated BMDMs **(Figure 1e).** Moreover, β-glucan-mediated dectin-1 signalling induces macrophage expression of early growth response (EGR) transcription factors 1-3^21^, which act as upstream transcriptional regulators of LPS-induced myeloid cell *F3* and *Serpine1* expression^22^. We found that β-glucan training prior to LPS stimulation caused significantly increased *EGR1* expression compared to LPS-stimulated BMDMs **(Figure 1b and g).**

**Figure 1:**
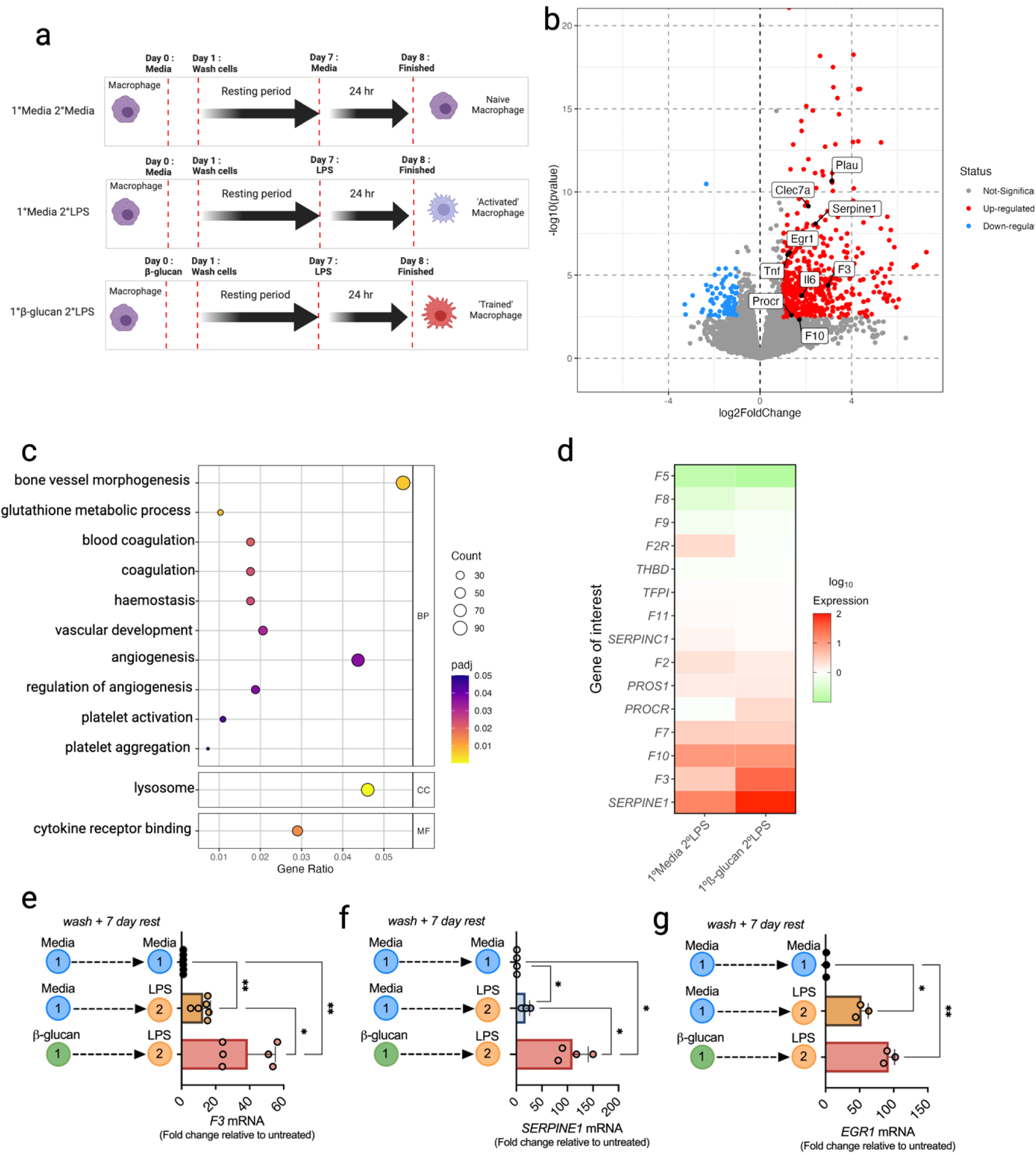
β-glucan-induced trained immunity upregulates coagulation-associated gene expression. **(a)** Schematic diagram outlining the protocol for inducing trained immunity in bone marrow-derived macrophages (BMDMs). Briefly, BMDMs were pre-treated with media or 100 µg/mL whole glucan particles and left for 24 hours before cells were washed 3 times with PBS and left to rest for 1 week. On day 7, cells were re-stimulated with 100 ng/mL LPS for 6 hours prior to RNA isolation. **(b)** Bulk RNA sequencing was performed on LPS-treated (n=3) and β-glucan-trained BMDMs re-stimulated with LPS (n=3), and differential expression of genes (DEG) analysis was performed between the groups. **(c)** Gene Ontology (GO) enrichment analysis of the gene sets significantly upregulated in LPS-restimulated β-glucan-trained BMDMs compared to LPS-treated BMDMs. (BP=biological processes; CC=cellular component; MF=molecular functions). **(d)** Heat map showing coagulation-associated gene expression profiles for LPS-treated (1°Media 2°LPS) and LPS-restimulated β-glucan-trained (1°β-glucan 2°LPS) BMDMs following RT-qPCR analysis. Data were normalised for *RPS18* mRNA levels, and results expressed as fold change compared to untreated BMDMs (1°Media 2°Media). RT-qPCR results for **(e)** *F3,* **(f)** *Serpine1* and **(g)** *EGR1* mRNA levels. A one-way ANOVA was used to determine statistical significance with *P≤0.05, **P≤0.01 for 6 independent experiments measured in duplicate.

### β-glucan-mediated training results in myeloid cell hypercoagulability

Based on these findings, we sought to determine the impact of β-glucan-mediated innate immune memory on macrophage procoagulant activity in plasma. Evaluation of β-glucan-trained BMDMs via a modified cell-based thrombin generation assay^23^ revealed that LPS-treated BMDMs previously trained with β-glucan possessed significantly increased procoagulant activity compared to LPS-treated BMDMs **(Figure 2a),** causing a 29-minute shortening of lagtime compared to naïve BMDMs, and a 22-minute reduction in lagtime compared to LPS-treated BMDMs **(Figure 2b).** Notably, no significant change in procoagulant activity was observed in BMDMs 24 hours after β-glucan exposure alone **(Supplementary Figure 3a)**, and *F3* gene expression was only minimally increased **(Supplementary Figure 3b)**. To confirm the enhanced procoagulant response in β-glucan-trained BMDMs was not elicited uniquely by LPS re-stimulation, β-glucan-trained BMDMs were re-stimulated with the TLR1/2 agonist PAM3CSK4. A similar increase in *F3* gene expression (31-fold) and decrease in lagtime (29-min) was observed in PAM3CSK4-restimulated trained BMDMs to that observed with LPS re-stimulated trained BMDMs. **(Supplementary Figure 4a-d)**.

**Figure 2:**
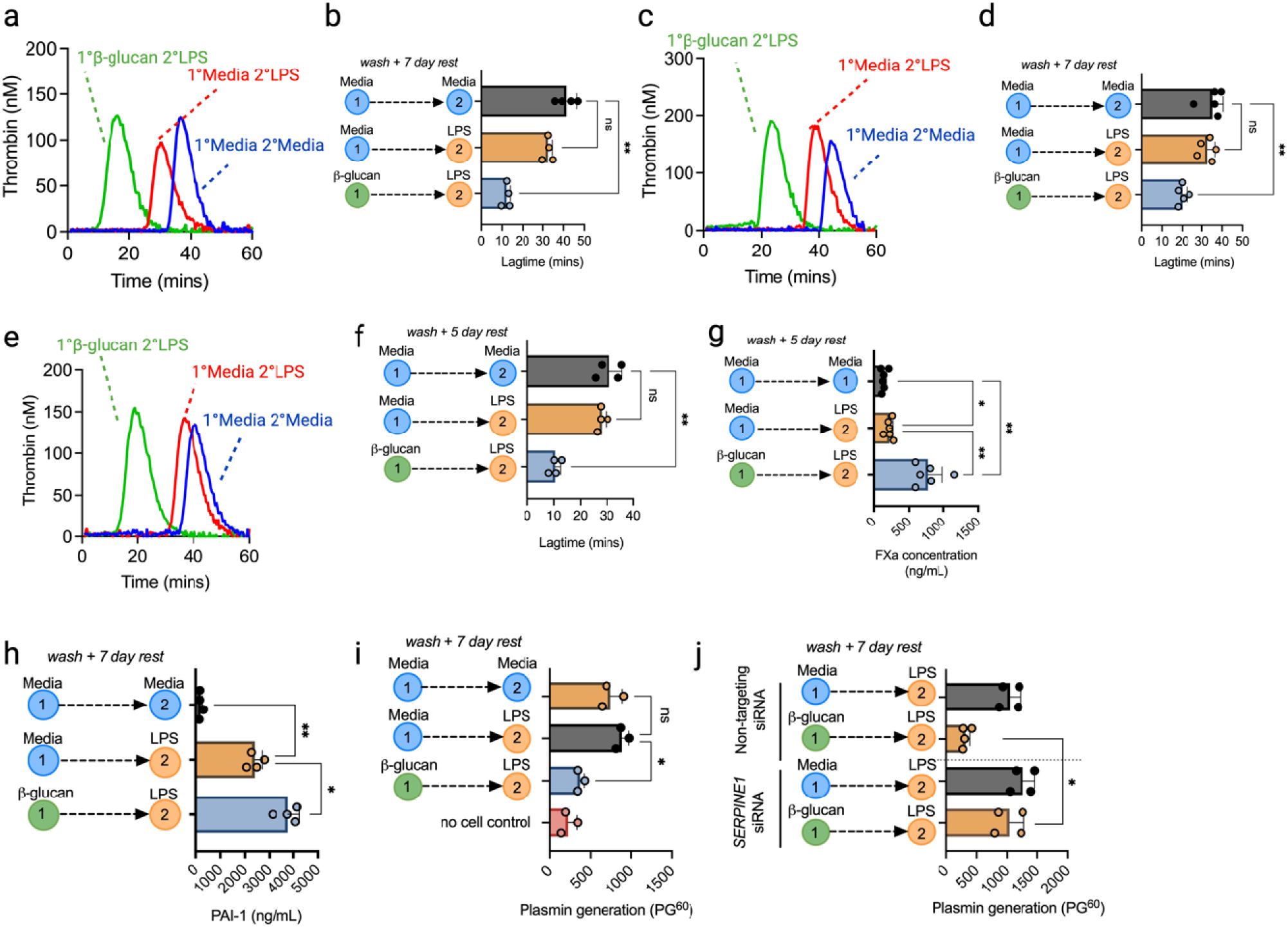
β-glucan-induced trained immunity enhances myeloid cell procoagulant activity. Cells were pre-treated with media or 100 µg/mL whole β-glucan particles and left for 24 hours before cells were washed 3 times with PBS and left to rest for 1 week. On day 7, cells were restimulated with 100 ng/mL LPS for 24 hours. A cell-based TGA was performed with cells as the sole source of TF activity in 80 µL FXII-deficient plasma, 20 µL MP reagent and 20 µL thrombin substrate added to initiate the reaction. **(a)** Thrombogram for TGA performed with BMDMs from each group and **(b)** the associated lag-times calculated. **(c)** Thrombogram for TGA performed in the presence of BMDM supernatants (80 µL) and **(d)** the associated lag-times that were determined. **(e)** Thrombogram for TGA performed with β-glucan-trained primary human monocytes and **(f)** the associated lag-times. **(g)** A FXa generation assay was performed in the presence of primary human monocytes incubated with 0.5 µg/mL FVIIa and 10 µg/mL FX for 30 minutes. FXa substrate was added to detect FXa generated due to TF activity. **(h)** PAI-1 protein levels measured by ELISA in β-glucan-trained BMDMs. **(i)** t-PA-mediated plasmin generation in the presence of β-glucan-trained BMDMs was measured using a plasmin-specific fluorogenic substrate, and the fluorescence reading after 60 mins (PG^60^) was determined. **(j)** BMDMs were transfected with *Serpine1* siRNA or non-targeting (NT) siRNA (both 20nM) before β-glucan training and LPS treatment. tPA-mediated plasmin generation in the presence of these cells was then assessed. A paired t-test or one-way ANOVA were used where appropriate to determine statistical significance with *P≤0.05, **P≤0.01 for 3-6 independent experiments measured in duplicate.

Supernatants from each treatment group were collected and assessed. Notably, a 12-minute shortening of lag time was observed in the presence of supernatants from re-stimulated β-glucan-trained BMDMs compared to supernatants from LPS-treated BMDMs, suggesting an increase in the presence of procoagulant microparticles in supernatants from β-glucan-trained BMDMs **(Figure 2c and d).** These data indicate that β-glucan-induced innate immune memory reduces the threshold for both LPS-induced TF procoagulant activity in macrophages and promotes procoagulant microparticle release.

Similarly, re-stimulated β-glucan-trained purified human monocytes shortened the lag time to thrombin generation by 20 minutes compared to naïve monocytes alone **(Figure 2e and f)**. FXa generation was also significantly enhanced in the presence of β-glucan-trained human monocytes compared to untreated or LPS-treated monocytes, confirming that the effect is TF-dependent **(Figure 2g)**. Recent studies have implicated T-lymphocytes in enabling trained immunity in human monocytes^24^. β-glucan-induced hypercoagulability was observed in both column-purified CD14^+^ monocytes and adherent PBMCs **(Supplementary Figure 5)**, suggesting that in contrast to their role in trained immunity, the presence of IFNγ-producing lymphocytes is not required for enhanced thrombin generation and procoagulant activity in β-glucan-trained monocytes.

### β-glucan-mediated training enhances LPS-mediated impairment of myeloid cell fibrinolytic activity

As re-stimulated β-glucan-trained BMDMs up-regulated *Serpine1* expression and doubled PAI-1 released compared to LPS-treated BMDMs **(Figure 1f and Figure 2h)**, we assessed the impact of re-stimulated β-glucan-trained BMDMs on macrophage regulation of fibrinolytic activity via altered plasmin generation. LPS re-stimulation of β-glucan-trained BMDMs exhibited significantly enhanced inhibition of plasmin generation compared to LPS-treated BMDMs **(Figure 2i).** Subsequent *Serpine1* inhibition by siRNA knockdown confirmed inhibition of plasmin generation resulted from increased PAI-1 production **(Figure 2j).** *Serpine1* gene expression was unchanged after 24 hours of β-glucan exposure and did not affect plasmin generation alone **(Supplementary Figure 3c-d)**. These data indicate that restimulated β-glucan-trained BMDMs possess a significantly lower threshold for procoagulant and antifibrinolytic activity than when directly stimulated by LPS alone.

### The protoporphyrin IX ring in free haem mediates trained hypercoagulability

We next sought to determine whether trained hypercoagulability could be induced by endogenous damage-associated molecular pattern (DAMP) mediators of trained immunity. Free haem has recently been shown to impart innate immune memory in myeloid cells^25^. Free haem treatment followed by LPS re-stimulation 7 days later caused a significant increase in TNFα production and metabolic reprogramming in both human monocytes and murine BMDMs, as previously reported **(Supplementary Figure 6a-b, e).** To first investigate the structural component(s) of free haem responsible for this response, individual haem structural components, protoporphyrin IX and Fe-NTA, were co-incubated with BMDMs, washed out, and re-stimulated with LPS 7 days later. Protoporphyrin IX pre-treatment caused a similar increase in TNFα production to that of free haem following LPS re-stimulation **(Supplementary Figure 6c)**, whereas Fe-NTA pre-treated BMDMs did not increase TNFα production after LPS re-stimulation **(Supplementary Figure 6d)**. Protoporphyrin-trained BMDMs also exhibited increased basal glycolysis compared to LPS-treated BMDMs, whereas FeNTA-treated BMDMs did not cause any change in basal glycolysis **(Supplementary Figure 6e)**. These results demonstrate that the protoporphyrin ring of haem induces metabolic reprogramming characteristic of trained myeloid cells.

*F3* gene expression in re-stimulated BMDMs exposed to haem one week prior was increased 9-fold relative to LPS-treated BMDMs **(Figure 3a).** Haem-trained BMDMs subsequently treated with LPS induced significantly shorter lag times to thrombin generation than BMDMs treated with LPS alone **(Figure 3b-c).** Interestingly, this effect largely depended on protoporphyrin IX, which shortened the lag time by 16 minutes compared to when naïve BMDMs were present **(Figure 3d-e).** Also, supernatants from re-stimulated haem-trained BMDMs caused a 15-minute shortening of lag time compared to LPS-treated BMDMs **(Figure 3f-g)**, suggesting haem-induced innate immune memory causes the enhanced release of procoagulant microparticles upon re-exposure to pro-inflammatory stimuli. Similarly, haem pre-treatment induced a lowered procoagulant threshold in purified human monocytes, resulting in a 13-minute shortening in lag time compared to LPS-treated BMDMs **(Figure 3h-i).**

**Figure 3:**
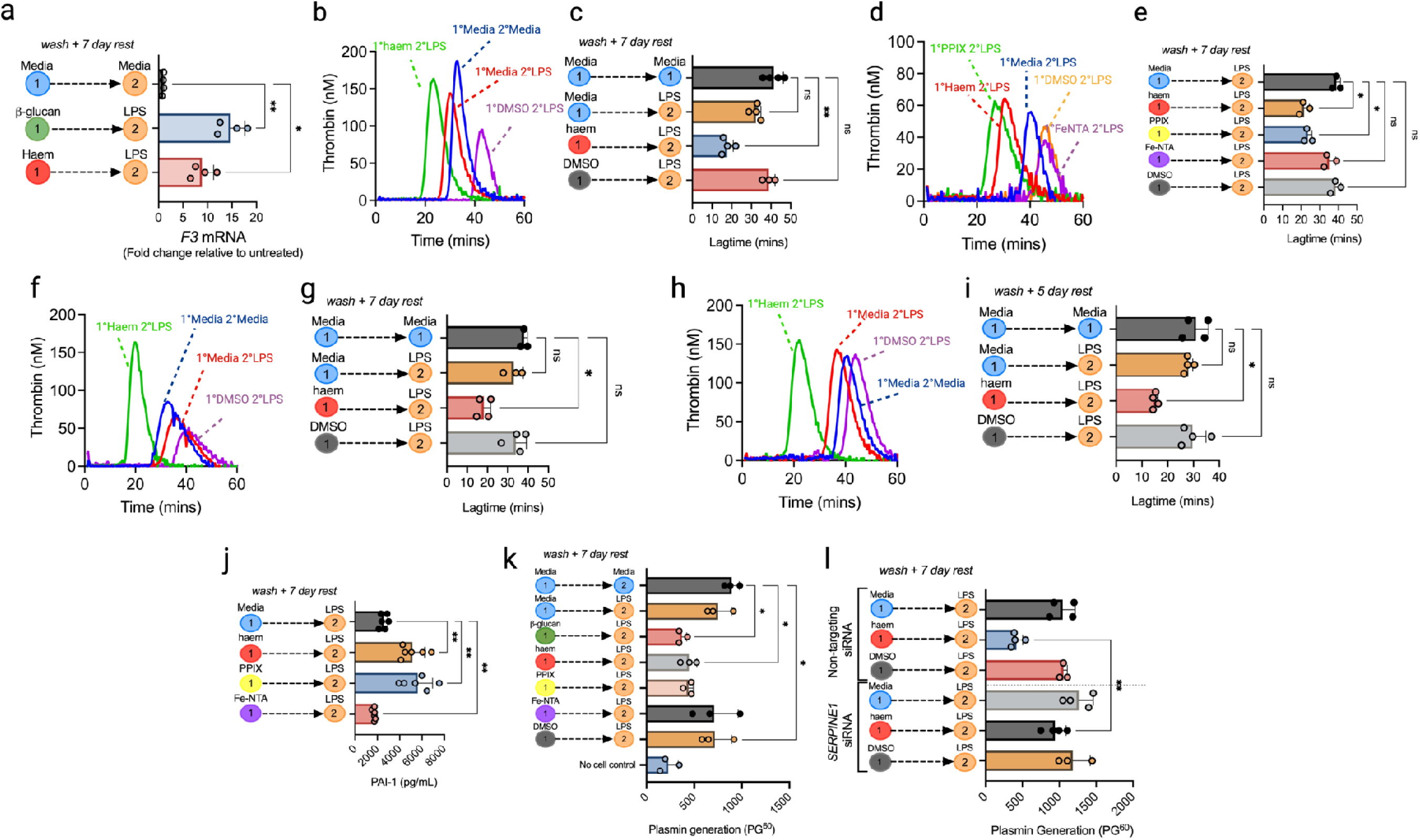
Free haem induces trained hypercoagulability in myeloid cells via the PPIX ring. BMDMs and human monocytes were trained with 50 µM haem, 50 µM protoporphyrin IX (PPIX), 50 µM Fe-NTA or DMSO vehicle control. After 24 hrs, cells were washed and rested for 7 days. Cells were then re-stimulated with 100 ng/mL LPS for 6 hrs for mRNA analysis and 24 hrs for protein analysis and functional assays. **(a)** *F3* mRNA levels in haem-trained BMDMs were determined by RT-qPCR. Data were normalised to *RPS18* mRNA levels, and results expressed as fold-change relative to untreated BMDMs (1°Media 2°Media). **(b)** Thrombogram for TGA performed in the presence of haem-trained BMDMs and **(c)** associated lag-times. **(d)** Thrombogram for TGA performed in the presence of PPIX or Fe-NTA trained BMDMs and **(e)** associated lag-times. **(f)** Thrombogram for TGA performed in the presence of haem-trained BMDM supernatants and **(g)** the associated lag-times. **(h)** Thrombogram for TGA performed in the presence of haem-trained human monocytes and **(i)** associated lag-times. **(j)** PAI-1 levels were measured by ELISA in haem-, PPIX- and Fe-NTA-trained BMDMs. **(k)** t-PA-mediated plasmin generation in the presence of haem-, PPIX- and Fe-NTA-trained BMDMs was measured using a plasmin-specific fluorogenic substrate, and the fluorescence reading after 60 mins (PG^60^) was determined. **(l)** BMDMs were transfected with *Serpine1* siRNA or non-targeting (NT) siRNA (both 20nM) before haem training and LPS treatment. tPA-mediated plasmin generation in the presence of treated BMDMs was then performed. A paired t-test or one-way ANOVA were used where appropriate to determine statistical significance with *P≤0.05, **P≤0.01 for 3-6 independent experiments measured in duplicate.

As with β-glucan-trained BMDMs, PAI-1 production was significantly increased in haem-trained BMDMs compared to LPS-treated BMDMs, which was similarly dependent on the protoporphyrin ring **(Figure 3j).** Both haem- and protoporphyrin-trained BMDMs reduced plasmin generation by approximately half of that generated by LPS-treated BMDMs, but there was no change in plasmin generation activity in Fe-NTA-treated BMDMs **(Figure 3k).** siRNA-mediated knockdown of *Serpine1* in haem-trained BMDMs resulted in recovery of plasmin generation comparable to LPS-treated BMDMs, confirming PAI-1-dependent inhibition of plasmin generation in re-stimulated haem-trained BMDMs **(Figure 3l).** These results demonstrate that trained hypercoagulability is a common feature of myeloid cell innate immune cell memory.

### Epigenetic and metabolic macrophage reprogramming are essential for trained hypercoagulability

Given the role of transcriptomic, epigenomic and metabolic reprogramming in enabling innate immune memory, we tested whether similar adaptations would be necessary for trained hypercoagulability. Firstly, we utilised inhibitors which target histone methylation and acetylation **(Figure 4a)**. In common with previous reports, the histone methyltransferase inhibitor, MTA, and the histone acetyltransferase inhibitor, EGCG, reduced macrophage TNFα release from re-stimulated β-glucan-trained BMDMs, whereas the histone demethylase inhibitor, PG, had no significant impact, in keeping with prior reports^26^ **(Figure 4b)**. Furthermore, *F3* expression was reduced 5-fold and 2-fold with MTA and EGCG pre-treatment in LPS re-stimulated, β-glucan-trained BMDMs **(Figure 4b).** PAI-1 production was also significantly impaired in MTA pre-treated trained BMDMs **(Figure 4b)**. These results suggest that post-translational histone modifications arising from β-glucan-induced innate immune memory are required for trained hypercoagulability and suggest a prominent role for histone methylation in the enhanced procoagulant activity of β-glucan-trained BMDMs. Innate immune memory also causes extensive rewiring of intracellular metabolic pathways, leading to enhanced macrophage glycolysis **(Supplementary Figure 1c)**. To examine the role of glycolysis in trained hypercoagulability, we utilised 2-deoxy-glucose (2-DG), a glucose analogue which acts as a competitive inhibitor of hexokinase. 2-DG treatment of β-glucan-trained BMDMs significantly prolonged thrombin generation, restoring lag time to that observed in the presence of untreated BMDMs **(Figure 4c and d),** demonstrating β-glucan-induced enhancement of macrophage glycolytic activity is necessary for trained hypercoagulability.

**Figure 4:**
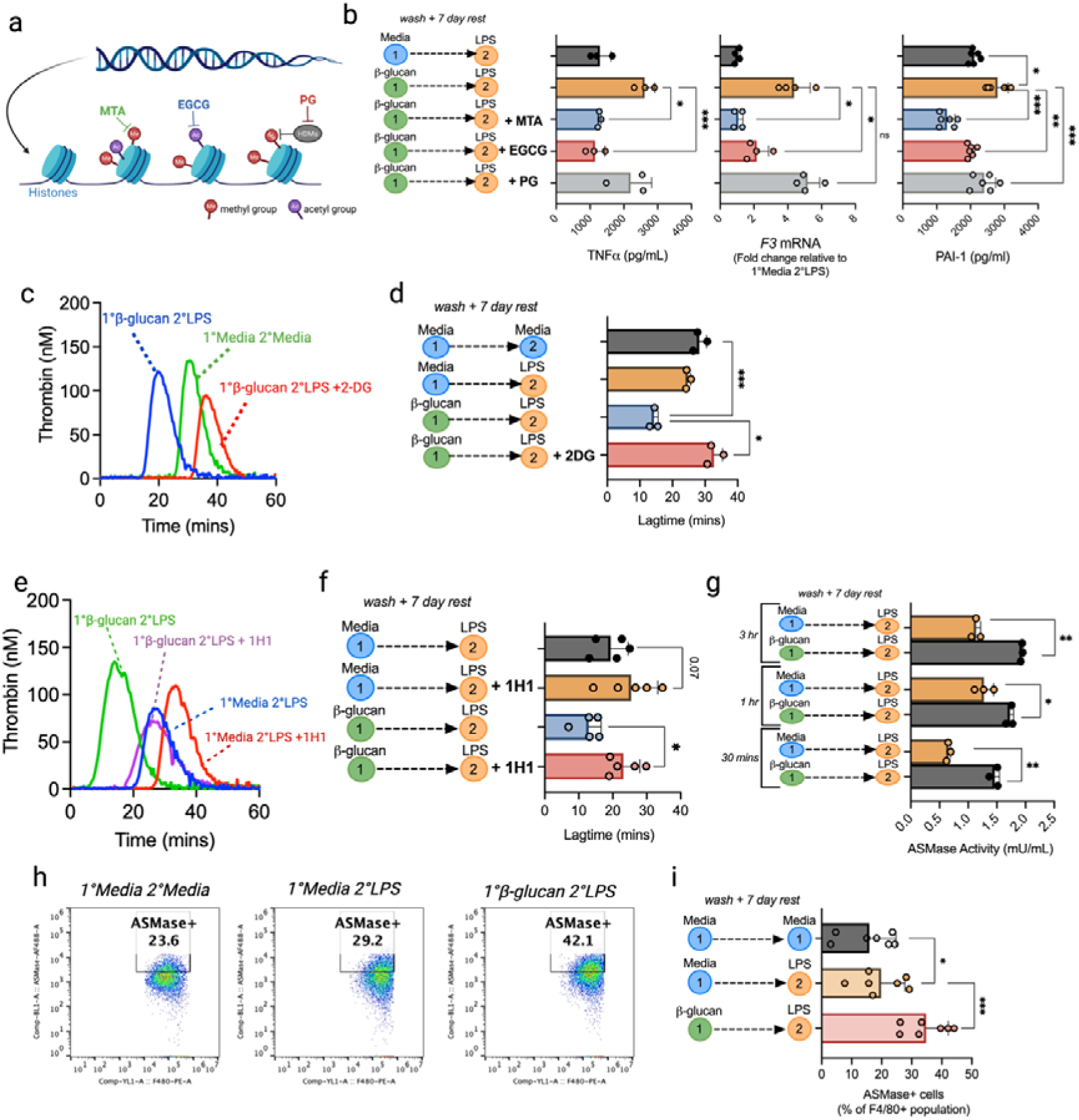
β-glucan induced trained hypercoagulability requires epigenetic and metabolic reprogramming. **(a)** Schematic diagram illustrating chemical epigenetic inhibitor targets. Methylthioadenosine (MTA) is a histone methyltransferase inhibitor, Epigallocatechin-3-gallate (EGCG) is a histone acetyltransferase inhibitor and paraglyine (PG) is a histone demethylase (HDM) inhibitor. **(b)** BMDMs were incubated with 1 mM MTA, 100µM EGCG or 12mM PG for 1 hr prior to training with 100 µg/mL β-glucan for 24 hr. Cells were washed 3 times and growth media supplemented with epigenetic inhibitors added for the rest period (100 µM MTA, 100 µM EGCG and 12 mM PG). On day 7, cells were re-stimulated with 100 ng/mL LPS. *TNFA* and *F3* mRNA levels in β-glucan-trained BMDMs were determined by RT-qPCR, and PAI-1 protein levels were determined by ELISA. **(c)** BMDMs were pre-treated for 3 hr with 10 mM 2-deoxy-D-glucose (2-DG), a glycolysis inhibitor, before β-glucan training protocol. After LPS re-stimulation on day 7, cells were assessed in a TGA to generate a thrombogram with **(d)** associated lag-times. **(e)** β-glucan-trained and LPS-stimulated BMDMs were incubated with an anti-TF antibody (1H1) for 1 hr prior to TGA analysis and **(f)** lag-time generation. **(g)** ASMase activity in trained BMDMs was measured by fluorogenic ASMase activity assay. Trained cells were restimulated with LPS for 30 mins, 1 hr or 3 hr prior to cell lysis. **(h-i)** ASMase cell surface expression on β-glucan-trained BMDMs was determined by flow cytometry in F4/80^+^ BMDM populations. A paired t-test or one-way ANOVA were used where appropriate to determine statistical significance with *P≤0.05, **P≤0.01, ***P≤0.001 for 3-7 independent experiments measured in duplicate.

### ASMase procoagulant activity is up-regulated in **β**-glucan-trained BMDMs

We next sought to investigate the mechanisms responsible for increased procoagulant activity observed in re-stimulated β-glucan-trained macrophages. The role of TF activity in mediating the enhanced procoagulant activity of restimulated β-glucan-trained BMDMs was confirmed using an anti-TF monoclonal antibody, which extended lag time to that of naïve BMDMs **(Figure 4e and f).** TF surface expression was increased in all treatment groups following LPS re-stimulation, however, despite the significant difference in procoagulant activity, there was surprisingly little difference in BMDM TF surface expression between LPS-treated BMDMs and LPS-treated BMDMs that had been previously exposed to β-glucan **(Supplementary Figure 7a)**. TF typically exists in an encrypted, non-coagulant state and additional mechanisms are required to decrypt TF into a procoagulant conformation. Phosphatidylserine (PS) externalisation is important for TF procoagulant activity and coagulation factor complex assembly necessary for thrombin generation and clot formation^27,28^. PS surface expression was investigated to examine whether PS externalisation was enhanced in β-glucan-trained BMDMs. LPS re-stimulation increased PS surface expression in all treatment groups, however, no significant difference was observed between trained BMDMs and LPS-treated cells **(Supplementary Figure 7b)**. Furthermore, changes in TF disulphide bond formation did not contribute to enhanced procoagulant activity in trained BMDMs, as treatment of macrophages with the PDI inhibitor rutin did not impact β-glucan-training mediated accelerated thrombin generation **(Supplementary Figure 7c-e)**. We next investigated how induction of innate immune memory may affect ASMase activity, which enhances TF procoagulant activity via TF decryption in macrophages^29^. Notably, β-glucan-trained BMDMs subsequently exposed to LPS had significantly elevated ASMase activity compared to BMDMs treated with LPS alone **(Figure 4g).** ASMase cell surface expression was also significantly increased on β-glucan-trained BMDMs compared to LPS-treated BMDMs **(Figure 4h and i)**. These results suggest that the enhanced TF activity observed in re-stimulated β-glucan-trained macrophages derives from enhanced TF decryption mediated by a training-mediated enhancement of cell surface ASMase activity.

### Induction of central trained immunity induces the generation of hypercoagulable myeloid cells

Systemic β-glucan training causes long-lasting epigenetic and metabolic alterations to haematopoietic progenitor cells in the BM that favour myelopoiesis^16^. To establish whether induction of central trained immunity would result in the generation of mature myeloid progeny with a lower threshold for increased procoagulant activity, mice were administered whole β-glucan particles by intraperitoneal injection and left for between 1-4 weeks **(Figure 5a)**. Splenic monocytes were then isolated from mice 1-4 weeks after β-glucan administration. To confirm that the isolated splenic monocytes exhibited characteristic features of trained immunity, TNFα production was measured before and after LPS re-stimulation. TNFα was not detected in monocytes before LPS re-stimulation, however, there was a significant increase in TNFα production in LPS re-stimulated monocytes from β-glucan-treated mice, with the most significant increase occurring in monocytes derived from mice that had received β-glucan 3-4 weeks prior **(Figure 5b).** IL-6 was also significantly increased in β-glucan-treated mice both before and after *ex vivo* LPS treatment in mice that had received β-glucan 3-4 weeks prior (**Figure 5c)**. Next, we compared the TF-mediated procoagulant activity of splenic monocytes from either saline- or β-glucan-administered mice. Splenic monocytes from mice treated with β-glucan 3-4 weeks previously exhibited a ∼6-minute shorter lag time than monocytes from saline-administered mice.

**Figure 5:**
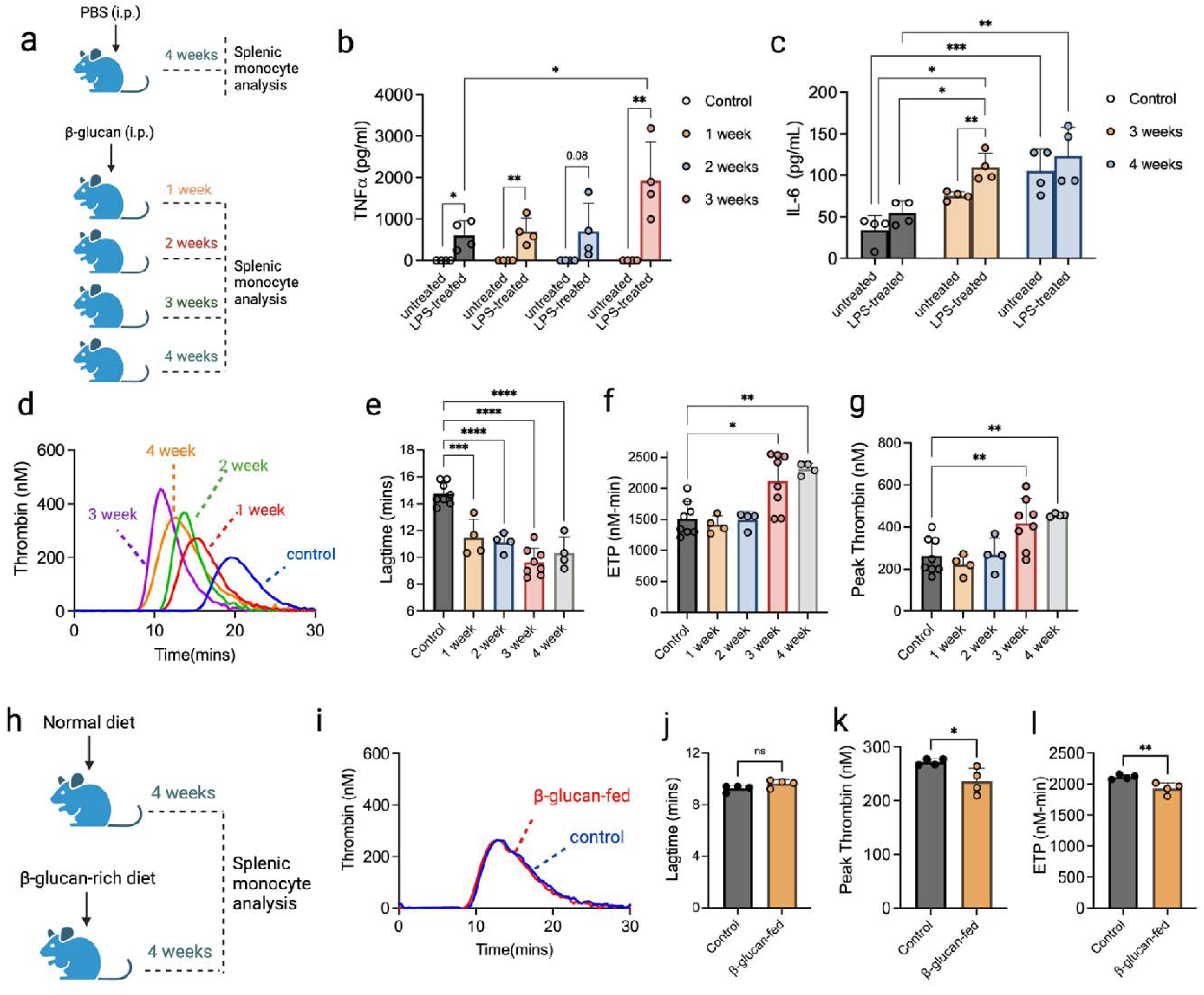
Splenic monocytes from β-glucan-administered mice display enhanced procoagulant activity weeks after administration. **(a)** A schematic depiction of the experimental outline to invoke trained immunity *in vivo*. Briefly, mice were injected with either PBS or whole glucan particles 1-4 weeks prior to sacrifice. CD115^+^ splenic monocyte population were then isolated. Splenic monocytes were left untreated or re-stimulated 100 ng/mL LPS ex vivo for 24 hr. **(b)** TNFα and **(c)** IL-6 levels released from splenic monocytes were determined by ELISA. **(d)** A TGA was performed in the presence of splenic monocytes isolated from mice administered β-glucan 1-4 weeks previously. **(e)** Lag-time, **(f)** ETP and **(g)** peak thrombin were also determined. **(h)** In parallel, mice were fed a standard diet or a diet supplemented with dietary β-glucan. Splenic monocytes were then isolated and assessed by TGA to give **(i)** thrombogram, **(j)** lag-time, **(k)** peak thrombin and **(l)** ETP. A paired t-test or one-way ANOVA were used where appropriate to determine statistical significance with *P≤0.05, **P≤0.01, ***P≤0.001, ***P≤0.0001 for 4-8 mice with samples measured in duplicate.

This phenomenon was observed in the presence and absence of *ex vivo* LPS re-stimulation **(Figure 5d and e, Supplementary Figure 8)**. Peak thrombin and ETP were also increased in the presence of monocytes isolated from mice treated with β-glucan 3 or 4 weeks previously compared to saline-administered mice, suggesting that central trained immunity *in vivo* causes the generation of monocytes with increased capacity for thrombin generation **(Figure 5f and g).** This procoagulant phenomenon plateaued at 3 weeks, with no further enhancement in TGA parameters after 4 weeks **(Figure 5d-g)**. Interestingly, the longer the interval between β-glucan administration and animal sacrifice, the more significant the increase in splenic monocyte procoagulant activity was observed. We also assessed whether dietary β-glucan had the same impact on splenic monocyte procoagulant activity **(Figure 5h)** but found no difference in splenic monocyte-mediated thrombin generation parameters between β-glucan-fed mice and control diet-fed mice **(Figure 5i-l).**

### Central trained immunity results in a hypercoagulable BM environment and increased plasma procoagulant activity

The detection of hypercoagulable myeloid cells that had been synthesised weeks after β-glucan administration suggested the development of a BM environment that favoured the generation of procoagulant mature myeloid cells. We next evaluated mouse bone marrow progenitor cell populations in saline-treated mice and mice administered β-glucan 3 weeks before sacrifice **(Figure 6a)**. As previously reported^16^, there was a significantly increased percentage of haematopoietic progenitor cells (LSK; Lin^−^cKit^+^Sca1^+^) in the BM of β-glucan-administered mice compared to saline-treated mice **(Figure 6b)** and within this population, a significant increase in myeloid-biased MPP3 (Flt3^−^CD48^+^CD150^−^LSK) cells **(Figure 6c, see Supplementary Figure 9 for gating strategy).** Notably, there was no significant increase in lymphoid-biased progenitors **(Supplementary Figure 10)**. As IL-1β enhances haematopoietic activity induced by trained immunity^9^, we measured IL-1β in the BM interstitial fluid of β-glucan-administered and saline-administered control mice. IL-1β was significantly increased in BM interstitial fluid isolated from β-glucan-administered mice **(Figure 6d)**, with a similar pattern observed for TNFα and IL-6 levels **(Figure 6e and f)**. Cytokine production from BMDMs isolated and cultured from β-glucan and saline-administered mice BM was also measured in the presence or absence of *ex vivo* LPS restimulation. TNF and IL-6 release was significantly increased from BMDMs isolated from β-glucan-administered mice upon LPS restimulation compared to BMDMs from saline-administered control mice **(Figure 6g and h)**. BMDMs generated from β-glucan-administered mice also supported enhanced thrombin generation compared to saline-treated mice both with and without *ex vivo* LPS treatment, as seen via reduced lagtime and enhanced ETP **(Figure 6i-k).**

**Figure 6:**
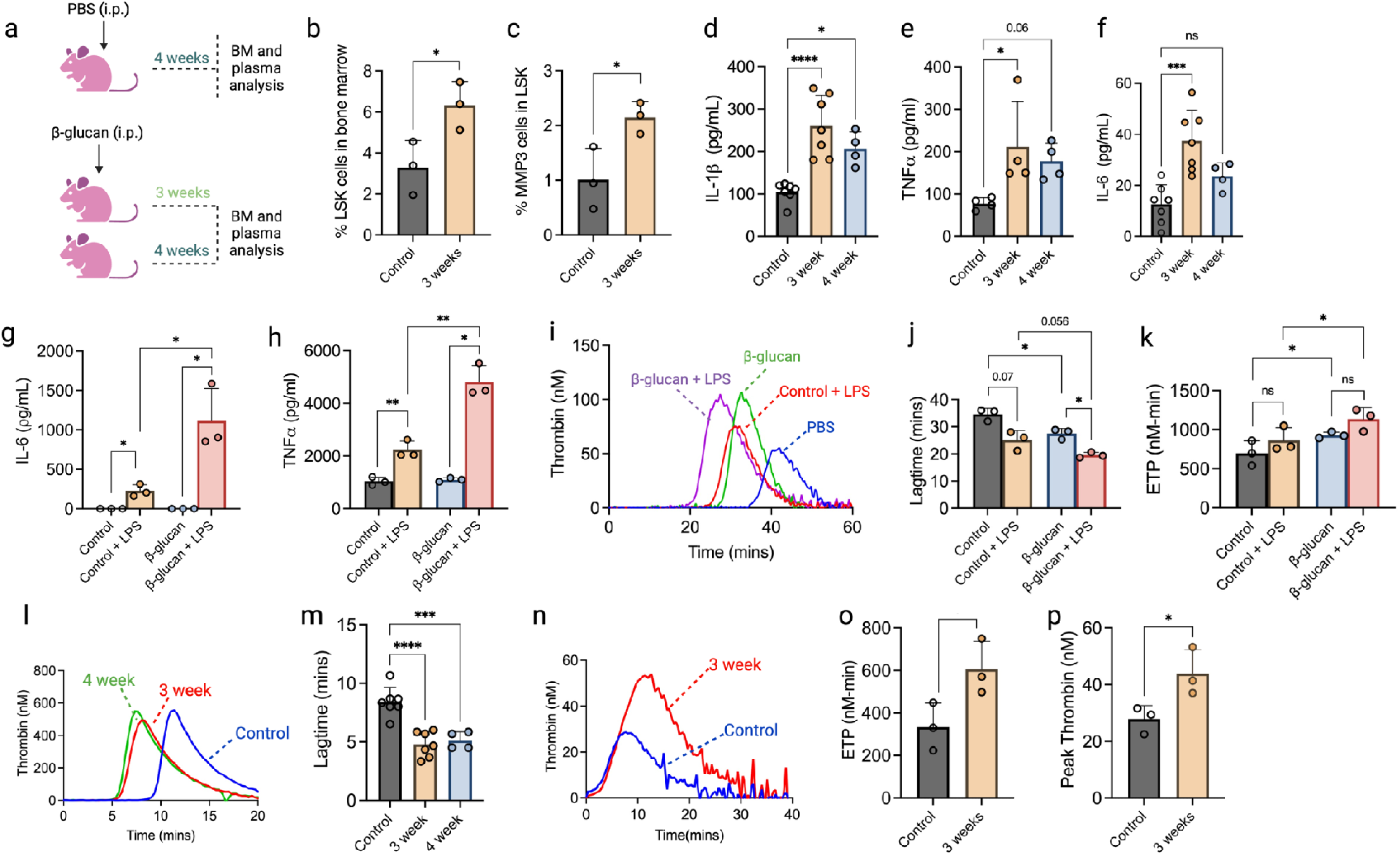
β-glucan-induced central trained immunity results in a BM environment with enhanced procoagulant activity. **(a)** Briefly, mice were injected with either PBS or whole glucan particles 1-4 weeks before sacrifice. Plasma was collected via the retro-orbital sinus. BM interstitial fluid and HSCs were harvested. BM haematopoietic stem and progenitor cell populations were analysed by flow cytometry for **(b)** % LSK^+^ cells in BM and **(c)** % MPP3 myeloid-biased progenitor cells in the LSK^+^ compartment. **(d)** IL-1β, **(e)** TNFα and **(f)** IL-6 cytokine levels were determined in BM interstitial fluid by ELISA. **(g)** IL-6 and **(h)** TNFα levels released from cultured BMDMs without or with *ex vivo* LPS re-stimulation were determined by ELISA. Cultured BMDMs without or with *ex vivo* LPS restimulation were included in a TGA to generate **(i)** thrombogram, **(j)** lag-time and **(k)** ETP values. **(l)** TGA was performed in the presence of BM interstitial fluid to give thrombogram and **(m)** associated lag-times **(n)** TGA analysis was performed with mouse platelet-poor plasma isolated from trained and control mice. Each well contained 20 µL plasma, 20 µL TBS and 20 µL FluCa thrombin substrate and TGA parameters **(o)** ETP and **(p)** peak thrombin values are shown. A paired t-test or one-way ANOVA were used where appropriate to determine statistical significance with *P≤0.05, **P≤0.01, ***P≤0.001, ***P≤0.0001 for 4-8 mice with samples measured in duplicate.

Given the increased pro-inflammatory cytokine production in β-glucan-administered mouse BM and the possibility of increased procoagulant microparticles, we assessed the procoagulant activity of cell-free BM interstitial fluid from β-glucan-administered and saline-administered mice. Notably, BM interstitial fluid from β-glucan-administered mice was able to drive quicker clot formation, shortening the lag time by ∼5 minutes compared to BM fluid from saline-administered mice, suggesting the increased presence of procoagulant microparticles within the mouse BM (**Figure 6l-m).** Finally, to assess whether prior β-glucan administration caused a systemic increase in plasma procoagulant activity, we assessed thrombin generation in platelet-poor plasma derived from β-glucan-administered and saline-treated control mice without the addition of an exogenous thrombin generation trigger. Plasma from mice administered β-glucan 3 weeks prior exhibited significantly higher peak thrombin and ETP than plasma from saline-administered control mice (**Figure 6n-p)**. Collectively, these results indicate that central trained immunity causes the adoption of a procoagulant BM microenvironment that results in the generation of myeloid cells with a lower procoagulant threshold and the release of procoagulant microparticles within the BM and the periphery.

## DISCUSSION

Immune cell adaptations in response to infection or inflammation are critical for host survival. Adaptive immunity leads to the development of antigen-specific lymphocytes that lay primed to respond to subsequent similar immune challenges. Innate immune memory, however, enables peripheral myeloid cells to respond to non-specific pathogens with enhanced vigour. This transformation is mediated by epigenetic and metabolic reprogramming that trains cells for subsequent immune responses to unrelated challenges. Although this feature remains beneficial in strengthening subsequent first-line defences to infection and malignancy, the ‘shortening of the inflammatory fuse’ has also been linked to the development of autoinflammatory diseases^30,31^. In this study, we demonstrate the prothrombotic consequences of inducing trained immunity in myeloid cells. While crucial for enhanced trained immune response, these adaptations inadvertently increase the propensity of trained myeloid cells to promote aberrant thrombin generation via dysregulated TF expression and decryption, combined with inhibition of macrophage profibrinolytic activity. Moreover, induction of central trained immunity by β−glucan administration in mice causes the generation of a pro-inflammatory BM environment defined not only by increased pro-inflammatory cytokine production, but also HSPC reprogramming that leads to the generation of myeloid cells with a lowered threshold for thrombin generation and increased procoagulant activity in BM interstitial fluid and plasma.

TF activity in myeloid cells rapidly increases upon exposure to pathogen- or damage-associated molecular patterns, but its expression typically subsides after 24 hours^32,33^. *F3* gene expression in activated myeloid cells is regulated by several transcription factors, including NF-κB, AP-1 family members and EGR1, that bind at the *F3* promoter and enhancer regions^34–36^. Our initial transcriptomic studies in BMDMs identified significantly increased *EGR1* expression when BMDMs were stimulated with β−glucan one week prior to LPS exposure, compared to BMDMs that were only exposed to LPS. In addition to regulating proinflammatory cytokine expression in response to infection or injury, EGR-1 is considered a master regulator transcription factor for many pro-inflammatory genes, including genes encoding TF and PAI-1^22,37^. Pertinently, previous studies have shown that *Candida albicans*-derived β-glucan stimulation of the dectin-1 receptor results in the activation of EGR-1, −2 and −3^21^. Given that β-glucan-mediated trained immunity is mediated through dectin-1 signalling^14^, EGR1 induction by β-glucan is likely central to subsequent activation of coagulation and antifibrinolytic gene programs in myeloid cells.

Durable innate immune memory in peripheral myeloid cells is made possible by the development of central trained immunity, in which a training agent increases BM IL-1β and granulocyte-macrophage colony-stimulating factor to promote myelopoiesis via HSPC changes in glucose metabolism and cholesterol biosynthesis pathways^16,38^. In assessing whether the observed β−glucan-mediated lowered threshold for coagulation was a transient phenomenon, or could be imparted long-term, we observed that in mice, isolated splenic monocyte procoagulant activity increased with time after β−glucan administration up to 3 weeks before plateauing at 4 weeks. Given the lifespan of monocytes in the periphery is 1-7 days, the observed hypercoagulability of these monocytes is in keeping with the synthesis of new centrally-trained peripheral myeloid cells with enhanced procoagulant, pro-inflammatory features. Interestingly, isolated monocytes from β−glucan-administered mice required *ex vivo* LPS stimulation to prompt enhanced TNF secretion, whereas IL-6 expression from isolated monocytes increased with the duration of time since β−glucan administration, with only a limited increase in response to additional LPS. Similarly, TF-driven thrombin generation mediated by isolated monocytes from β−glucan-administered mice was only minimally enhanced by additional LPS. This suggests that β−glucan-induced central trained immunity, in addition to skewing BM activity towards myelopoiesis, also causes the generation of monocytes with a constitutively lower threshold for procoagulant activity. Although the duration of trained hypercoagulability is currently unknown, it is possible it could persist for longer than the 3-4 weeks observed in our study. Previous studies have shown that the epigenetic and metabolic reprogramming associated with trained immunity *in vivo* typically persist for weeks to months,^18,39,40^ however, in human studies, particularly in relation to BCG vaccination-mediated trained immunity, hyper-responsive innate immune cells can persist for years^41^. Remarkably, some studies have also suggested that trained immunity may be inherited from prior generations. One study showed that the progeny of mice that survived an infection of *C. albicans* had enhanced survival when subjected to endotoxin challenge, and this protection was conferred via epigenetic and transcriptional changes in the myeloid progenitor cell BM cell populations^42^. Additionally, in humans, the beneficial effects of the BCG vaccine in newborns were enhanced when the mother had also received the BCG vaccine^43^.

Similarly, we observed significantly increased procoagulant activity in BM interstitial fluid from mice administered β−glucan >3 weeks before sacrifice. This suggests the increased release of procoagulant microparticles into the BM microenvironment during central training. Although the cellular sources of procoagulant microparticles in the BM have not yet been determined, previous studies have shown that HSCs in the BM release TF-containing extracellular vesicles, which contribute to thrombus formation following laser-induced vessel wall injury^44^. Additionally, other studies have shown that depletion of TF expression on HSCs protects against endotoxin challenge by reducing coagulation activation^45,46^. Increased plasma procoagulant activity was also observed, although whether this is a consequence of TF-containing extracellular vesicles generated in the BM entering the bloodstream, or whether TF-containing extracellular vesicles were instead more readily released from newly synthesised monocytes recently generated in the BM or other BM-derived cells is not yet known.

Importantly, trained hypercoagulability in myeloid cells was not limited to β-glucan training, as we also observed the same similar procoagulant and anti-fibrinolytic response in cells trained with free haem. Haem was recently shown to also induce myeloid cell-trained immunity in monocytes and macrophages. This is mediated by epigenetic reprogramming, specifically through H3K27ac, which was enriched at hundreds of genomic loci in haem-trained BMDMs^11^. Increased circulating free haem via haemolysis arises in a number of inflammatory diseases, but is particularly important in the context of sickle cell disease (SCD)^47^. Circulating immune cells from individuals with SCD are often characterised by a poorly understood heightened pro-inflammatory state, which contributes to vaso-occlusive crisis in affected individuals^48,49^. Sickle RBCs are prone to haemolysis, releasing RBC intracellular components into the intravascular space to activate the surrounding endothelium and circulating leukocytes^50^. In particular, haem released from lysed RBCs can induce pro-inflammatory cytokine release from monocytes, further amplifying the pro-inflammatory conditions within the vasculature of SCD patients, but the molecular basis for this is not well-understood^51,52^. Haemoglobin released from lysed RBCs contains a reactive ferric protoporphyrin-IX group, which can trigger vascular and organ dysfunction and cause adverse clinical effects, including renal failure, immune dysregulation, atherosclerosis, vascular injury and endothelial dysfunction^53,54^. Interestingly, free haem has also been shown to contribute to TF-dependent thrombin generation in a mouse model of SCD^55^. Despite this, hemopexin, which neutralises free haem, only partially attenuated thrombin generation in sickle cell mice, suggesting the presence of additional contributory mechanisms^55^. Our data suggests that increased free haem may also promote both peripheral and central trained immunity to further exacerbate the risk of coagulopathy in SCD patients. These data could also provide mechanistic insights into the pathophysiology of other haemolytic conditions, such as thrombotic thrombocytopenic purpura and haemolytic uremic syndrome, which are characterised by increased labile haem in the circulation and a significantly increased risk of thrombotic complications^56,57^.

Collectively, these findings raise important questions about the potential contribution of immune cell adaptation in response to inflammatory stimuli and the mechanistic basis for the long-term prothrombotic risk observed in chronic or autoimmune inflammatory disease patients. Furthermore, the prothrombotic nature of trained myeloid cells warrants further consideration in relation to approaches to modulate trained immunity for therapeutic purposes^26,58–62^.

## ACKNOWLEDGEMENTS

Grant support for RJSP is provided by Science Foundation Ireland (21/FFP-A/8859), The National Children’s Research Centre (C/18/3) and Health Research Board (ILPPOR-2022-060).

## AUTHOR CONTRIBUTIONS

A.M.R. and R.J.S.P. devised the study and experimental strategy and wrote the manuscript. S.M., G.L., P.A.K., A.E.L., T.A.J.R., H.C-L., E.H.G., E.A.D., C.M., J.S.O’D, A.M.C., L.A.J.O’N., F.J.S., performed experiments and analysed data. All authors reviewed and contributed to the final version of the manuscript.

## MATERIALS AND METHODS

### Generation of BM-derived macrophages (BMDMs)

Female wild-type C57/BL6 mice aged between 6-12 weeks were euthanised by cervical dislocation, and hind legs were dissected, removed, and stored in high glucose DMEM (Merck) supplemented with 10% foetal bovine serum (ThermoFisher Scientific) 1U/mL penicillin and 0.1mg/mL streptomycin (Merck). Macrophages were isolated as previously described ^63,64^. Isolated macrophages were re-suspended in supplemented DMEM containing 20% M-CSF derived from L929 cells and incubated for six days, receiving 1mL M-CSF and 4mL fresh growth media on day 3. After six days, macrophages were detached using PBS containing 5mM EDTA (Merck) and plated for experiments in complete DMEM containing 10% M-CSF.

### Isolation and culture of human monocytes

Anonymised healthy donor buffy coats were obtained from the Irish Blood Transfusion Service, St. James’ Hospital, Dublin and monocytes were isolated as previously described^65^. Briefly, PBMCs were isolated from buffy coats by layering blood over Histopaque-1077 gradient (Merck) and centrifuging at 400xg for 25 minutes with acceleration and brake set to 1. Monocytes where then isolated using a Percoll (Merck) density centrifugation with 48.5% Percoll, 41.5% sterile water (Sigma) and 0.16 M NaCl (Merck) at 600 x g for 15 min with acceleration and brake set to 1. Monocytes were plated in serum-free RPMI containing 10 mM sodium pyruvate for 1 hr prior to changing media to complete RPMI supplemented with 10% human serum (from human male AB plasma, Merck), 1 U/mL penicillin and 0.1 mg/mL streptomycin. Alternatively, a pure monocyte population was obtained from PBMCs using CD14+ Microbeads (Miltenyi Biotec).

### Innate immune cell training in BMDMs

BMDMs were plated at 5 × 10^5^ cells/well in 12 well plates for RNA isolation, or 5 × 10^4^ cells/mL for TGA, PGA and ELISAs in 96 well plates with complete DMEM and 10% M-CSF and left overnight to adhere. Cells were treated with either 100 ng/mL LPS (Enzo Life Sciences), 100 µg/mL β-glucan (Invivogen), 50 µM Hemin (Merck), 50 µM protophorphyrin (Merck) or 50 µM Fe-NTA. Treated cells were left for 24 hr before aspirating media and washing 3 times with pre-warmed PBS. DMEM containing 10% M-CSF was added to wells and cells left for a 7-day rest period. During this period, M-CSF and media were changed every 3 days. On day 7, all treatment wells (excluding control wells) were incubated with 100 ng/mL LPS, before being washed with PBS and either lysed for gene expression analysis, or included in other assays.

### Innate immune cell training in human monocytes

Human monocytes were plated at 2 × 10^5^ cells/well in 96 well plates and cultured for 2-3 days in RPMI supplemented with 10% human serum (from human male AB plasma, Merck), 1 U/mL penicillin and 0.1 mg/mL streptomycin. Cells were then treated with 100 ng/mL LPS (Enzo Life Sciences, 100 µg/mL β-glucan (Invivogen) or 10-50 µM Hemin (Merck) for 24 hrs. Following incubation, cells were gently washed with PBS three times, and growth media was replaced. Cells were left to rest for 5 days with media replaced on day 3. Cells were re-stimulated with 100 ng/mL LPS on day 5. On the day of assay, cell plates were centrifuged at 1200 rpm for 5 min prior to removal of supernatants and gently washing cells with PBS. After this, downstream assays were performed.

### Myeloid cell-dependent thrombin generation assay (TGA)

Myeloid cell-dependent TGA was performed as previously described^23^. Briefly, following cell stimulations, supernatants were removed, and cells were washed once with PBS. 20µL MP-reagent was added to 80µL factor XII (FXII)-deficient plasma (Protylix). To examine the procoagulant activity of each cell releasate, 80µL of supernatants were added to plasma and MP reagent. The assay was initiated using 20µL FluCa-kit. Fluorescence was quantified using Thrombinoscope software on a Fluoroskan fluorometer.

### Murine plasma TGA

Thrombin generation potential in murine plasma was determined without exogenous TF trigger. Assay performed with wells containing 20 µL murine plasma, 20 µL TBS and 20 µL FluCa-kit to initiate the reaction. The temperature for the assay run was lowered to 33°C on the fluorometer to overcome the increased coagulation inhibitor activity in mouse plasma compared to human plasma, and calibrator activity on the Thrombinoscope software was set to 20% higher to account for a lower rate of conversion of the thrombin substrate by murine thrombin. These optimisations were based on previous literature^66^. Fluorescence was quantified using Thrombinoscope software on a Fluoroskan fluorometer.

### Myeloid cell-dependent plasmin generation assay (PGA)

Myeloid cell-dependent PGA was performed as previously described^23^. Briefly, following cell stimulations, macrophages were washed with PBS and then incubated with 100µL serum-free DMEM supplemented with 1U/mL penicillin, 0.1mg/mL streptomycin, 200ng/mL t-PA and 400nM Glu-plasminogen (Protylix) for 1.5 hours. Supernatants were then transferred to a black flat-bottomed 96-well plate. 20µL Boc-Glu-Lys-Lys fluorometric substrate was added to each well to enable plasmin detection (final concentration, 1260µM in TBS containing 34mM CaCl_2_; Bachem). Assays were performed with each treatment group in duplicate per assay and at least biological triplicates, with a negative control well containing only serum-free DMEM. Fluorescence was quantified on a Fluoroskan Ascent Fluorometer.

### FXa generation assay

Cell surface TF activity was measured as the ability of human monocyte monolayers to support FXa generation in the presence of activated factor VIIa (FVIIa) and CaCl_2_, as previously described^67,68^. Monocytes were seeded at 1 × 10^6^ cells/mL in 96 well plates. Briefly, cells were washed twice with buffer A (10 nM HEPES, 0.15 M NaCl, 4 mM KCl, 11 mM glucose, pH 7.5) after which cells were incubated with 100 µL Buffer B (buffer A containing 5 mg/mL BSA and 5 mM CaCl_2_) containing 0.5 µg/mL FVIIa (Protylix) and 10 µg/mL FX (Protylix) for 30 minutes. After this time, 25 µL was collected from each well and added to 50 µL Buffer C (50 mM Tris-HCl, 0.15 M NaCl, pH 7.5, with 1 mg/mL BSA and 25 mM EDTA). 50 µL of this mixture was then transferred to a new 96-well plate, and 50 µL of 1.25 mg/mL substrate was added to each well. The plate was then read at 405 nm for 1 hour taking a reading every 30 seconds. The amount of FXa generated in the sample wells was calculated by interpolating from the V_max_ of a FXa standard curve starting at 1000 ng/mL using serial dilutions of purified human FXa (Protylix).

### mRNA isolation and RT-qPCR

5 × 10^5^ cells/mL were lysed using 350µL lysis buffer and total RNA isolated according to manufacturer’s instructions (Ambion PureLink RNA isolation kit). Gene expression was determined by performing RT-qPCR in duplicate with Power Up SYBR green master mix (Thermofisher) using a 7500 Fast system (Applied Biosystems). Relative-fold changes in mRNA expression were calculated using the cycle threshold (C_T_) and normalised to the *RPS18* housekeeping gene. Primer sequences are listed in **Supplementary Table 1**.

### ELISAs

PAI-1, TNFα, IL-6 and IL-1β levels released by treated macrophages were determined using DuoSet ELISA kits (R&D Systems) per the manufacturer’s protocol.

### Macrophage siRNA transfection

SMARTpool siRNA guides were designed by Horizon Discovery and chosen using the siGENOME siRNA search tool. siRNAs (20nM) were transfected using Lipofectamine RNAiMax (Invitrogen). Cells were seeded at 2.5 × 10^5^ cells per well on 24-well plates. On the day of transfection, media was removed and replaced with 400µL of OptiMEM (Gibco) containing 5% M-CSF. siRNAs were transfected into cells at a concentration of 20nM using Lipofectamine RNAiMax (Invitrogen). Untransfected controls containing OptiMEM were also included in each experiment. The following day, media was removed and replaced with DMEM containing 10% M-CSF derived from L-929 cells. 24 hours following transfection, cells were treated as required for downstream assays.

### In vivo trained immunity model

Male C57BL/6J-OlaHsd mice aged 8-12 weeks were used. All experiments were performed under the approval of the Health Products Regulatory Authority, Ireland and Trinity College Dublin Animal Research Ethics Committee (AE19136/P148). Trained immunity was induced in mice with a single intraperitoneal (i.p.) injection of either 200 μL of PBS as a control or 200 μL of whole glucan particles (Invivogen) resuspended in PBS at 1 mg/mL. At week 1, week 2, week 3 or week 4 post-injection, whole blood was collected from animals via the retro-orbital sinus into 3.2% sodium citrate-coated tubes and plasma was obtained by centrifuging at 2000 rpm at 4°C for 15 mins. Mice were euthanised by CO_2_ inhalation, and spleens and legs harvested. To assess the effect of dietary β-glucan, mice were fed a diet supplemented with dietary fibre whole glucan particle (Kerry Food group). After 4 weeks, mice were euthanised by CO_2_ inhalation and spleens were harvested as before.

### Isolation of splenic monocytes and macrophages

Spleens were homogenised and passed through a 70 μm cell strainer (BD) with RPMI 1640 media supplemented with 10% FBS, 1 U/mL penicillin and 0.1 mg/mL streptomycin to obtain a single cell suspension for each sample. Each cell suspension was centrifuged at 1200 rpm for 5 min, and the supernatant was discarded. Splenocytes were then resuspended in 1 mL of red cell lysis buffer for 2 min before washing with 10 mL of complete RPMI media. Each sample was centrifuged again at 1200 rpm for 5 min), the supernatant was discarded, and cells were resuspended in 5 mL complete RPMI. Monocytes were then isolated using anti-CD115 microbeads (Miltenyi Biotec) according to manufacturer protocol. Following isolation, cells were centrifuged and resuspended in RPMI supplemented with 10% FBS, 1 U/mL penicillin, and 0.1 mg/mL streptomycin before plating at the required cell densities. Cells were plated at 5 × 10^4^ cells/well on 96 well plates for functional assays. Cells were left to adhere overnight, and fresh growth media was added the following day. Cells were then treated with 100 ng/mL LPS for 24 hours for functional assays.

### Collection of BM interstitial fluid

To collect BM interstitial fluid, mouse femurs were flushed with 500 µL cold PBS, and the fluid was centrifuged at 500 x g for 5 mins at 4°C. The BM fluid was then collected, transferred to new Eppendorf tubes, and stored at −80°C for further analysis.

### Cell mito stress test

OCR and ECAR were analysed in BMDMs using the Seahorse XFe96 Analyser (Agilent). The Cell Mito Stress Test assay was performed according to manufacturer instructions. 5 × 10^4^ cells/well were seeded onto Seahorse XF96 cell culture microplate and left to adhere overnight. BMDMs were then trained as described. BMDMs were then re-stimulated with 100 ng/mL LPS for 6 hr before performing a Cell Mito Stress Test. Supernatants were removed and replaced with assay medium (Seahorse XF DMEM (Agilent) supplemented with 1 mM pyruvate, 2 mM glutamine and 10 mM glucose (Merck)). The assay was performed with sequentially injected 1 μM oligomycin, 1 μM FCCP, and 0.5 μM rotenone/antimycin A. All measurements were performed with 8 technical replicates per experiment. Once the assay was completed, data was normalised to total protein per well determined by a BCA Protein Assay Kit (Thermo Fisher Scientific). Alterations in cellular metabolism were assessed by changes in OCR and ECAR using the Wave Desktop Software.

### Flow cytometry

Following each treatment, macrophages were incubated in PBS containing 5 mM EDTA for 5 min to remove cells from 12 well plates. Cells were washed with 1 mL PBS and stained with Live/Dead^TM^ dye (Invitrogen) for 30 mins at 4°C in the dark. Cells were left on ice for 30 min in the dark, after which FACS tubes were centrifuged at 1200 rpm for 5 min and supernatants decanted. Cells were then incubated with an anti-CD16/CD32 monoclonal antibody to block FC receptors (Invitrogen). Cells were stained with murine anti-F4/80 antigen PE conjugated antibody (Invitrogen), murine anti-TF 1H1 antibody (Genentech) and ASMase Polyclonal antibody (Invitrogen) on ice in the dark for 1 hr. The secondary antibody for TF staining was goat anti-rat IgG Alexa Fluoro 568 (Invitrogen) while the secondary for ASMase staining was goat anti-rabbit IgG Alexa Fluoro 488 (Invitrogen). Cells were washed and resuspended in 300 µL FACS Buffer and analysed using the BD FACs Canto flow cytometer for TF expression and Annexin V and the Attune NxT Flow Cytometer for F4/80 and ASMase expression.

### Measurement of phosphatidylserine (PS) exposure on BMDMs

PS expression was assessed using Annexin V and PI staining. For Annexin V staining, all washes, incubations and measurements were performed in Annexin V binding buffer (ThermoFisher Scientific). PI staining was performed in PBS. BMDMs were incubated with 1 μg/mL Annexin V-FITC (ThermoFisher Scientific). After 5 mins, cells were washed and resuspended in 200 μL of binding buffer. PI was added directly to FACS tubes (1 μg/mL) immediately before acquiring the samples on the BD FACs Canto flow cytometer.

### Measurement of acid-sphingomyelinase (ASMase) activity

Trained macrophages were restimulated with 100 ng/mL LPS on day 7 for 30 mins, 1 hour or 3 hours before cells were lysed. ASMase activity in the cell lysates was then measured using an ASMase activity assay kit (Echelon Biosciences) according to the manufacturer’s instructions.

### Flow cytometry analysis of HSPCs populations

To analyse HSPC populations in mouse BM after in vivo induction of trained immunity, isolated BM cells were isolated and cKit^+^ cells were enriched for using cKit Microbeads (Miltenyi). Isolated cKit^+^ HSPCs were resuspended in flow buffer; PBS-1X (Gibco) supplemented with 2% heat-inactivated FBS (Gibco). A flow cytometry staining protocol was applied to all BM samples as previously described^17^. Cells were stained with fixable viability stain ZombieAqua (Biolegend) at 1:1000 for 15 minutes. The following antibodies were then used for staining Lin^-^ c-Kit^+^ Sca^+^ cells (LKS), hematopoietic stem cells (HSCs) and multipotent progenitors (MPPs): anti-Ter-119 (clone TER-119, Biolegend), anti-CD5 (clone 53-7.3, Biolegend), anti-CD8a (clone 53-6.7, Biolegend), anti-B220+ (clone RA3-6B3, Biolegend), anti-Ly6G/C+ (clone RB6-8C5, Biolegend), all conjugated to APC-Cy7 were added 1:200 for 25 minutes. Other extracellular antibodies for anti-c-Kit-APC (clone 2B8, Biolegend), anti-Sca-1-PE-Cy7 (clone D7, eBioscience), antiCD150– PE (clone TC15-12F, eBioscience), anti-CD48-PerCP-eFluor710 (clone HM48-1, BD Bioscience), anti-CD34-FITC (clone RAM34, eBioscience), anti-Flt3-BV421 (clone A2F10.1, Biolegend) were added at 1:50 and incubated at 4°C for 25 minutes. All cells were washed with flow buffer and resuspended with IC Fixation Buffer (Invitrogen). Compensation controls were obtained after staining UltraComp eBeads Compensation Beads (Invitrogen) with the appropriate antibodies. Absolute counts were obtained using Precision Count Beads (Biolegend). Cells were acquired on the BD flow cytometer Fortessa with FACSDiva software. Data analysis was performed using FlowJo software. See **Supplementary Figure 9** for the gating strategy used.

### RNA sequencing and analysis

Splenic monocytes were isolated from control and β-glucan injected mice and restimulated *ex vivo* with 100 ng/mL LPS for 6 hours. Supernatants were removed, cells were washed once with PBS, and cells were lysed with lysis buffer (Ambion PureLink^TM^ RNA isolation kit) containing 1% 2-mercaptoethanol. Total RNA was isolated according to the manufacturer’s instructions. RNA samples were purified using RNeasy PowerClean Pro Cleanup Kit (Qiagen). Sample QC, RNA library preparation and QC were performed in-house by Novogene, Cambridge. RNA sequencing was performed using Illumina PE150 technology. Raw data from RNA sequencing was then bioinformatically processed in-house by Novogene. Briefly, this included quality control of reads where read removed if they included adaptor contamination, >10% of bases were resolved as uncertain (N), and where >50% of bases had a base quality < 5. All samples had >97% of bases with a Q Phred values greater than 20, and over 93% of bases with a Q Phred value greater than 30. With these filtered samples and reads, reads were aligned to the mouse GRCm38 reference using HISAT2^69^. Using the counts of reads to each mapped gene as input, differential gene expression analysis was carried out with the R package DESeq2^70^ using the recommended pipeline by its authors. The enrichment analysis was carried out using the clusterProfiler software^71^.

Volcano plot visualisation of log_2_-Fold Change and log_10_-transformed p-values, gene ontology enrichment visualisation, and heat map visualisation of gene expression profiles between LPS-restimulated β-glucan-trained BMDMs compared to LPS-treated BMDMs was carried out using the statistical computing language R version 4.3.2^72^ and the ggplot2 R package^73^

