## Supplementary Materials for "Trained Immunity Causes Myeloid Cell Hypercoagulability"

**Rehill et al SUPPLEMENTARY DATA**

**SUPPLEMENTARY FIGURE LEGENDS:**

**Supplementary Figure 1: β-glucan trained BMDMs have increased proinflammatory cytokine production and increased glycolysis.** BMDMs were pre-treated with media or 100 µg/mL whole glucan particle, left for 24 hr before cells were washed 3 times with PBS and left to rest for 1 week. On day 7, cells were restimulated with 100 ng/mL LPS. **(a)** TNFα production from BMDMs was measured by ELISA. **(b-c)** XF Seahorse Mito Stress Test was performed on β-glucan trained cells to determine extracellular acidification rate (ECAR) following the sequential addition of oligomycin, FCCP and Rotenone/antimycin. Results are shown as ECAR basal glycolysis. A paired t-test was used to determine statistical significance with *P≤0.05 for 3-4 independent experiments measured in duplicate.

**Supplementary Figure 2: RNAseq analysis of LPS-treated BMDMs.** RNA sequencing was performed on LPS-treated (n=3) and untreated BMDMs (n=3). **(a)** Differential expression of genes (DEGs) analysis carried out between the groups. **(b)** Gene Ontology (GO) enrichment analysis of the gene sets which are significantly upregulated in LPS-treated BMDMs compared to untreated BMDMs. (BP=biological processes; MF= molecular functions).

**Supplementary Figure 3: Single treatment with β-glucan does not induce TF or PAI-1 activity** BMDMs were treated with media or β-glucan for 24 hours before performing a TGA and RNA isolation. **(a)** TGA was performed with these BMDMs to generate lag-times. **(b)** *F3* and **(c)** *Serpine1* expression were determined by RT-qPCR. **(d)** t-PA-mediated plasmin generation in the presence of BMDMs was measured using a plasmin-specific fluorogenic substrate, and the fluorescence reading after 60 mins (PG^60^) was determined. A paired t-test or one-way ANOVA was used where appropriate to determine statistical significance with *P≤0.05, **P≤0.01, ****P≤0.0001 for 3-4 independent experiments measured in duplicate.

**Supplementary Figure 4: PAM3CSK4 restimulation following β-glucan-mediated training also increases BMDM hypercoagulability.** BMDMs were pre-treated with media or 100 µg/mL β-glucan, left for 24 hr before cells were washed 3 times with PBS and left to rest for 1 week. On day 7, cells were restimulated with 50 µg/mL PAM3CSK4. **(a)** *F3* mRNA levels were determined by RT-qPCR. **(b)** TGA was performed with PAM3CSK4 restimulated β-glucan primed-BMDMs and **(c)** associated lag-time and **(d)** peak thrombin determined. A paired t-test or one-way ANOVA was used where appropriate to determine statistical significance with *P≤0.05, **P≤0.01, ****P≤0.0001 for 3-4 independent experiments measured in duplicate.

**Supplementary Figure 5: IFNγ-producing lymphocytes are not required for enhanced procoagulant activity in β-glucan-trained monocytes.** Peripheral blood mononuclear cells (PBMCs) were isolated from healthy donor buffy coats by gradient centrifugation using histopaque-1077. PBMCs were then plated in presence of sodium pyruvate to encourage adherence. Alternatively, purified monocytes were isolated from PBMC using CD14^+^ positive selection microbeads. PBMC and CD14+ monocyte populations were primed with 100 µg/mL β-glucan, left for 24 hr before cells were washed 3 times with PBS and left to rest for 5 days. On day 5, cells were restimulated with 100 ng/mL LPS for 24 hr before TGA analysis was performed. Representative thrombograms for **(a)** PBMC monocytes and **(b)** CD14^+^ purified monocytes and associated parameters **(c)** lag-times, **(d)** ETP and **(e)** peak thrombin were determined. A one-way ANOVA was used to determine statistical significance with *P≤0.05 and ***P≤0.001 for 3-4 independent experiments measured in duplicate.

**Supplementary Figure 6: Free haem induces trained immunity in myeloid cells through enhanced pro-inflammatory cytokine production and increased glycolysis.** BMDMs and human monocytes were trained with 100 µg/mL β-glucan, 50 µM haem, 50 µM protoporphyrin IX (PPIX), 50 µM Fe-NTA or DMSO vehicle control. After 24 hrs, cells were washed and rested for 7 days. Cells were then restimulated with 100 ng/mL LPS. TNFα levels were measured by ELISA in **(a)** haem-trained human monocytes, **(b)** haem-trained BMDMs, **(c)** PPIX-trained BMDMs and **(d)** Fe-NTA-trained BMDMs. **(e)** XF Seahorse Mito Stress Test was performed on free haem- , PPIX- and Fe-NTA-trained BMDMs to determine ECAR following sequential addition of oligomycin, FCCP and Rotenone/antimycin. A paired t-test was used to determine statistical significance with *P≤0.05 and **P≤0.01 for 3-4 independent experiments measured in duplicate.

**Supplementary Figure 7: β-glucan-mediated trained immunity does not increase TF expression, PS externalisation or protein disulfide isomerase (PDI).** BMDMs were pre-treated with media or 100 µg/mL β-glucan, left for 24 hr before cells were washed 3 times with PBS and left to rest for 1 week. On day 7, cells were either left untreated or restimulated with 100 ng/mL LPS. **(a)** TF surface expression on live cells was measured by flow cytometry, **(b)** Phosphatidylserine exposure was measured by examining Annexin V binding by flow cytometry. Rutin, a potent inhibitor of PDI, was incubated (100 µM) with β-glucan and LPS-treated BMDMs for 1 h before TGA analysis. Representative thrombograms for **(c)** LPS-treated BMDMs, **(d)** β-glucan-trained BMDMs and **(e)** associated lag-times were determined. A paired t-test was used to determine statistical significance with *P≤0.05 and **P≤0.01 for 3-4 independent experiments measured in duplicate.

**Supplementary Figure 8: *Ex vivo* LPS stimulation of splenic monocytes from β-glucan administered mice is not required for enhanced procoagulant activity.** β-glucan mice were injected with either PBS or whole glucan particles 1-4 weeks before sacrifice. CD115^+^ splenic monocyte population were then isolated. Splenic monocytes were restimulated 100 ng/mL LPS *ex vivo* for 24 hr. A TGA was performed in the presence of LPS-treated splenic monocytes to generate **(a)** thrombogram, **(b)** lag-time and **(c)** ETP values. A One-Way ANOVA was used to determine statistical significance with *P≤0.05, **P≤0.01 and ****P≤0.0001 for 4-8 mice with samples measured in duplicate.

**Supplementary Figure 9: Gating strategy for flow cytometry analysis of haematopoietic stem and progenitor cell (HSPCs) populations.** Flow cytometry gating strategy for haematopoietic stem cells (HSCs) and multipotent progenitor (MPP) populations. LSK (Lin^−^cKit^+^Sca1^+^), Common myeloid progenitor (CMP; Lin^-^CD127^-^ckit^+^Sca-1^-^CD34^+^CD16/32^-^)

granulocyte macrophage progenitors (GMP; Lin^-^CD127^-^ckit^+^Sca-1^-^CD34^+^CD16/32^+^), common lymphoid progenitors (CLP; Lin^-^CD127^+^Ckit^+^Sca1^+^), myeloid biased MPP3 (Sca-1^+^ckit^+^CD48^+^CD150^−^Flt3^-^ and lymphoid biased MPP4 (Sca-1^+^ckit^+^CD48^+^CD150^−^Flt3^+^).

**Supplementary Figure 10: No significant change in lymphoid progenitor populations in β-glucan administered mice.** Mice were injected with either PBS or whole glucan particles 1-4 weeks prior to sacrifice. BM haematopoietic stem and progenitor cell populations were analysed by flow cytometry for **(a)** % MPP4 lymphoid-biased progenitor cells in the LSK^+^ compartment, and (b) % CMP, (c) % GMP, **(d)** % CLP in the bone marrow. An unpaired t-test was used to determine statistical significance for 3 biological replicates.

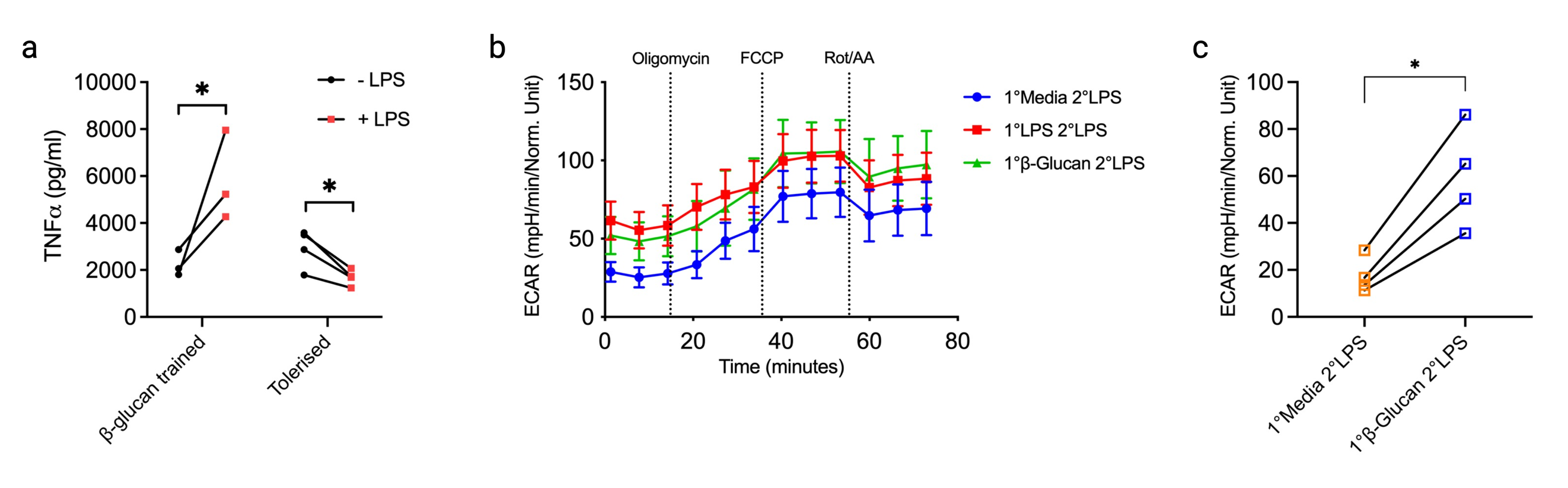

**Supplementary Figure 1**

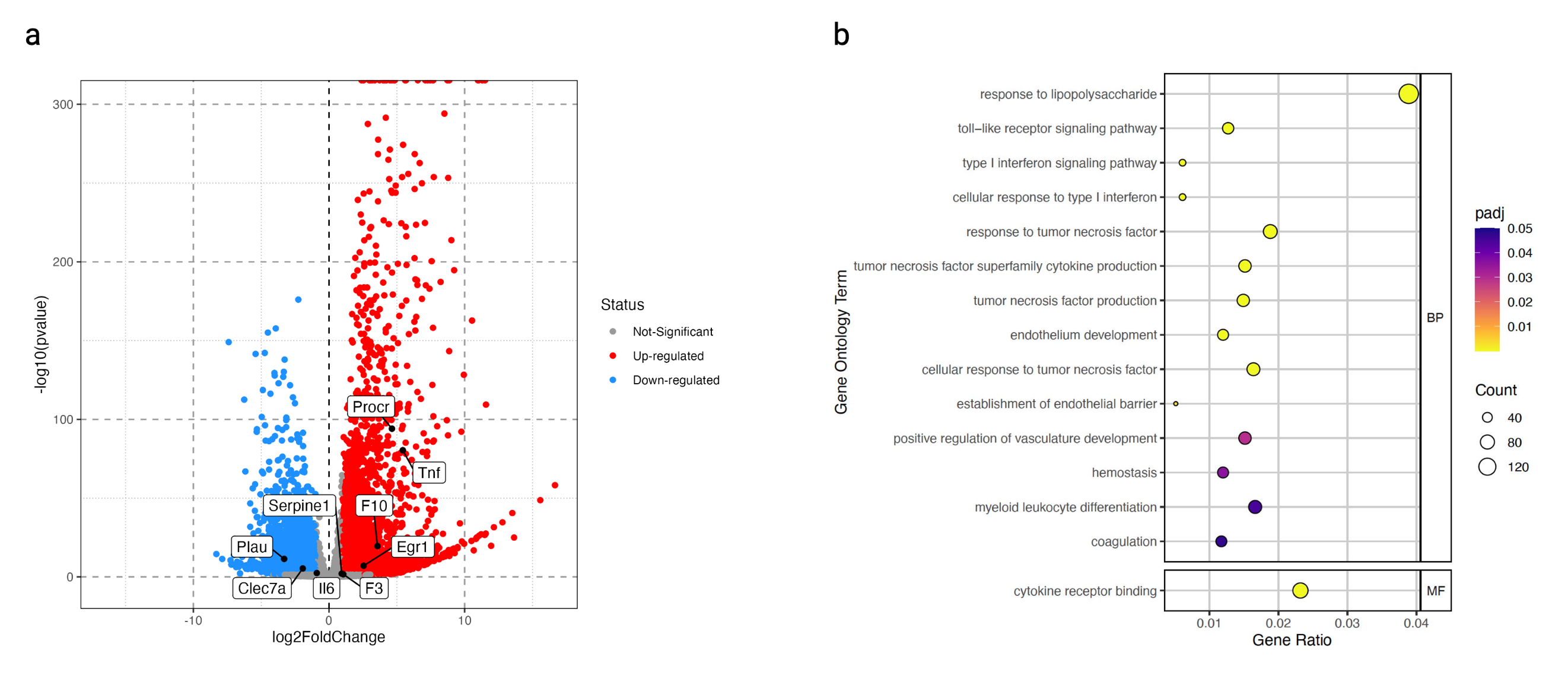

**Supplementary Figure 2**

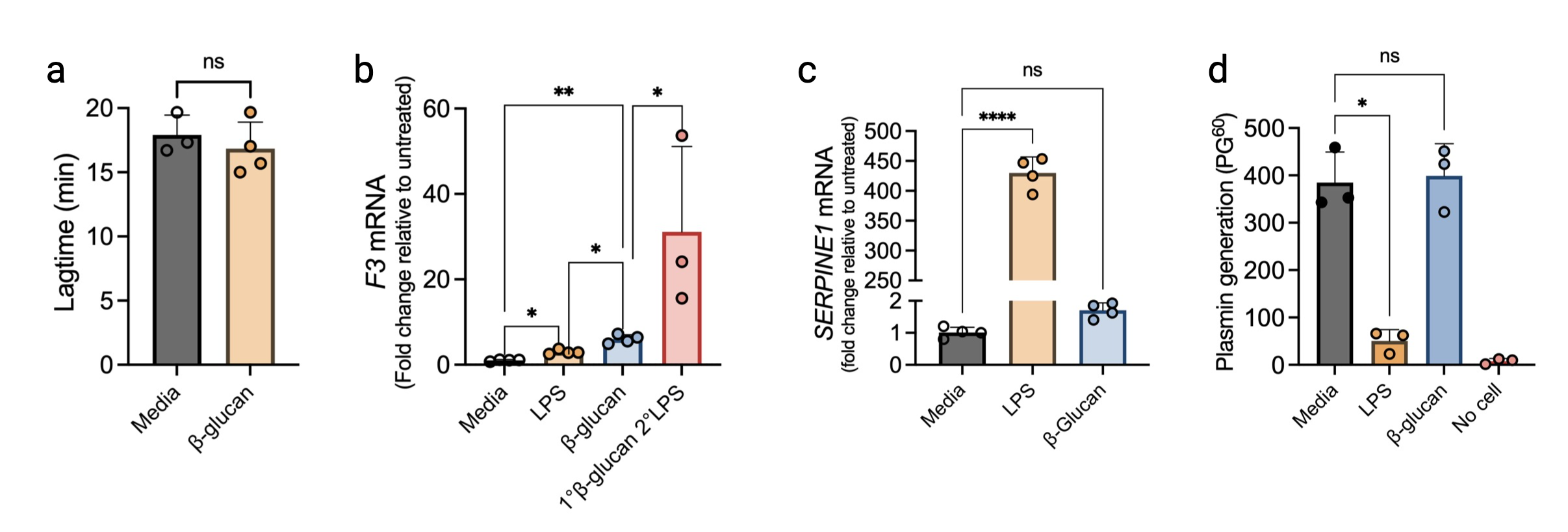

**Supplementary Figure 3**

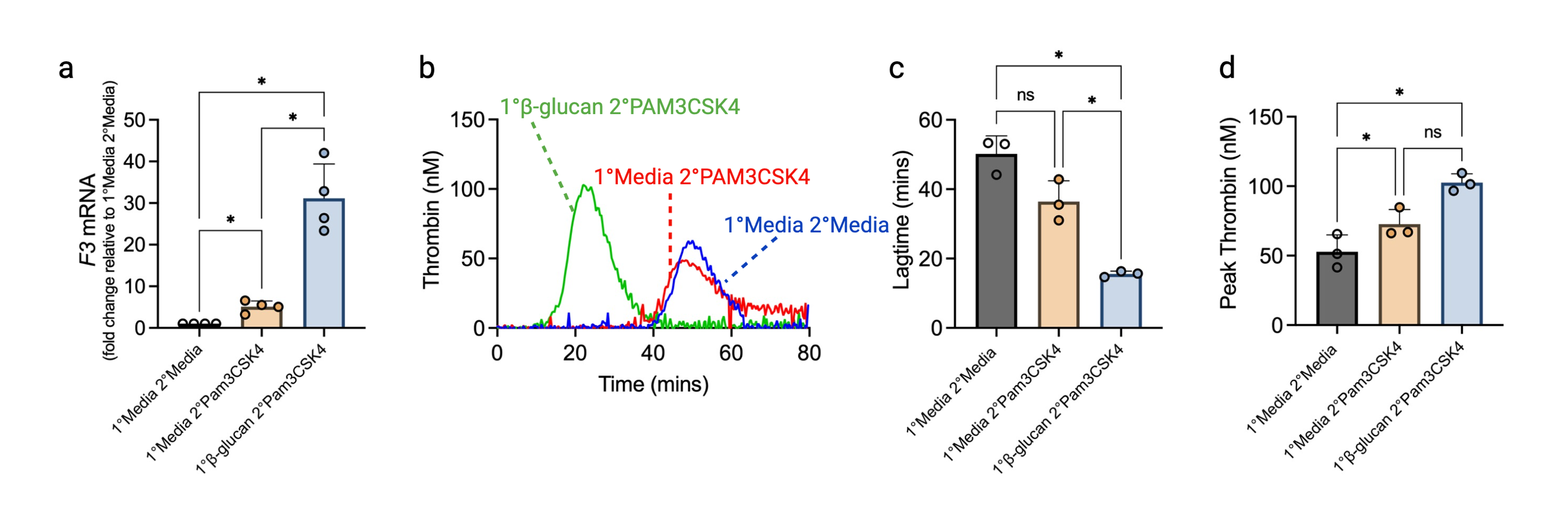

**Supplementary Figure 4**

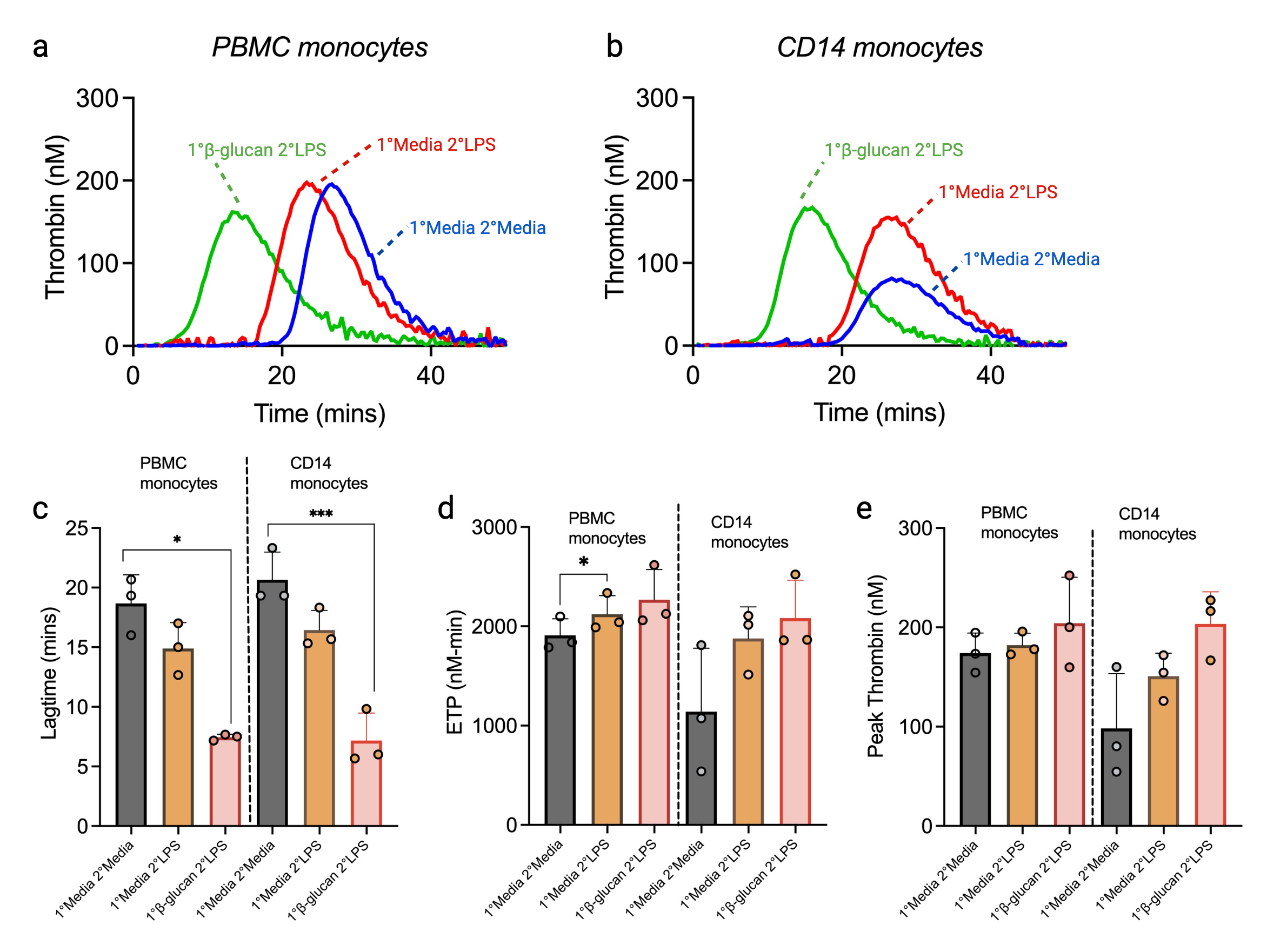

**Supplementary Figure 5**

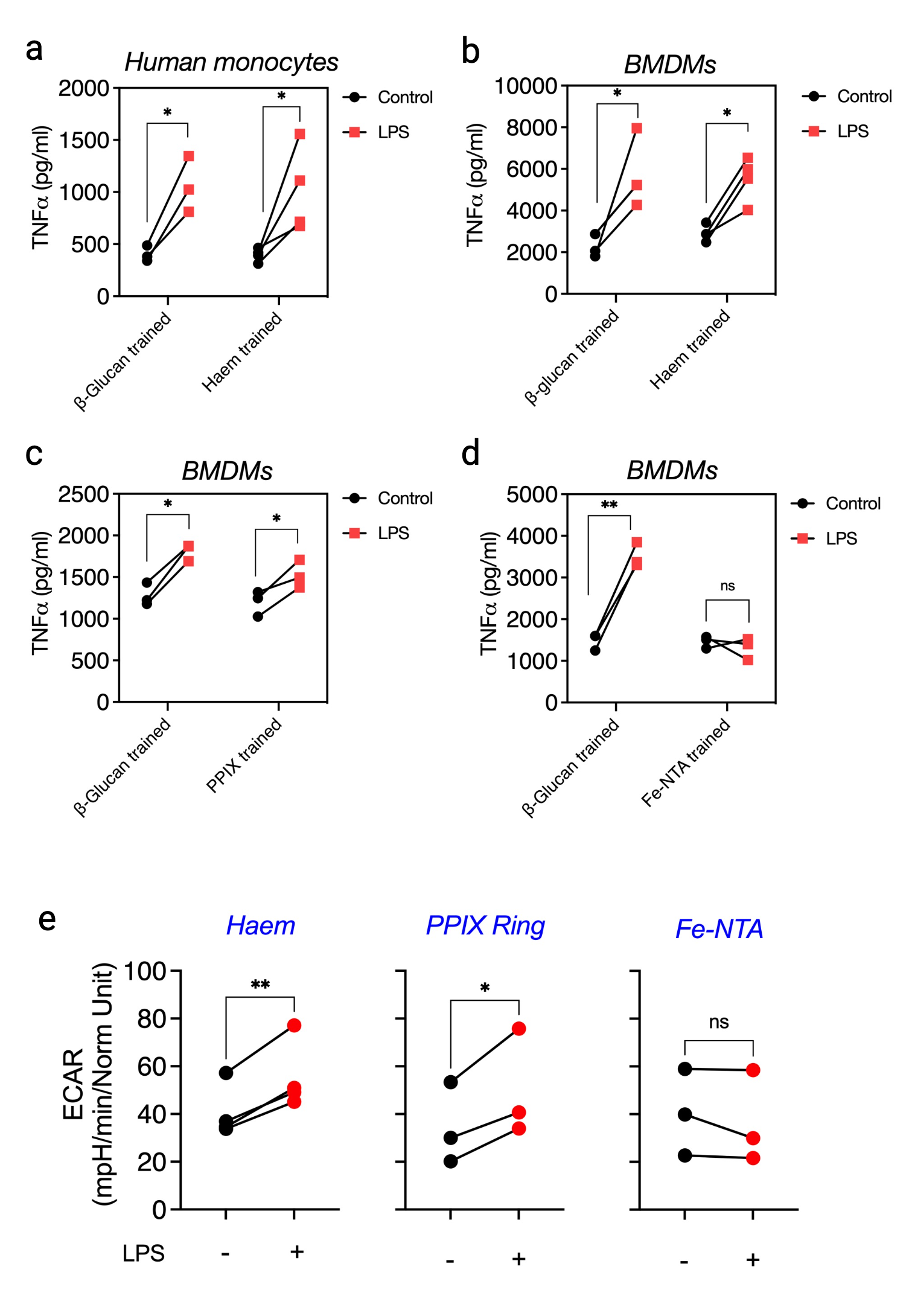

**Supplementary Figure 6**

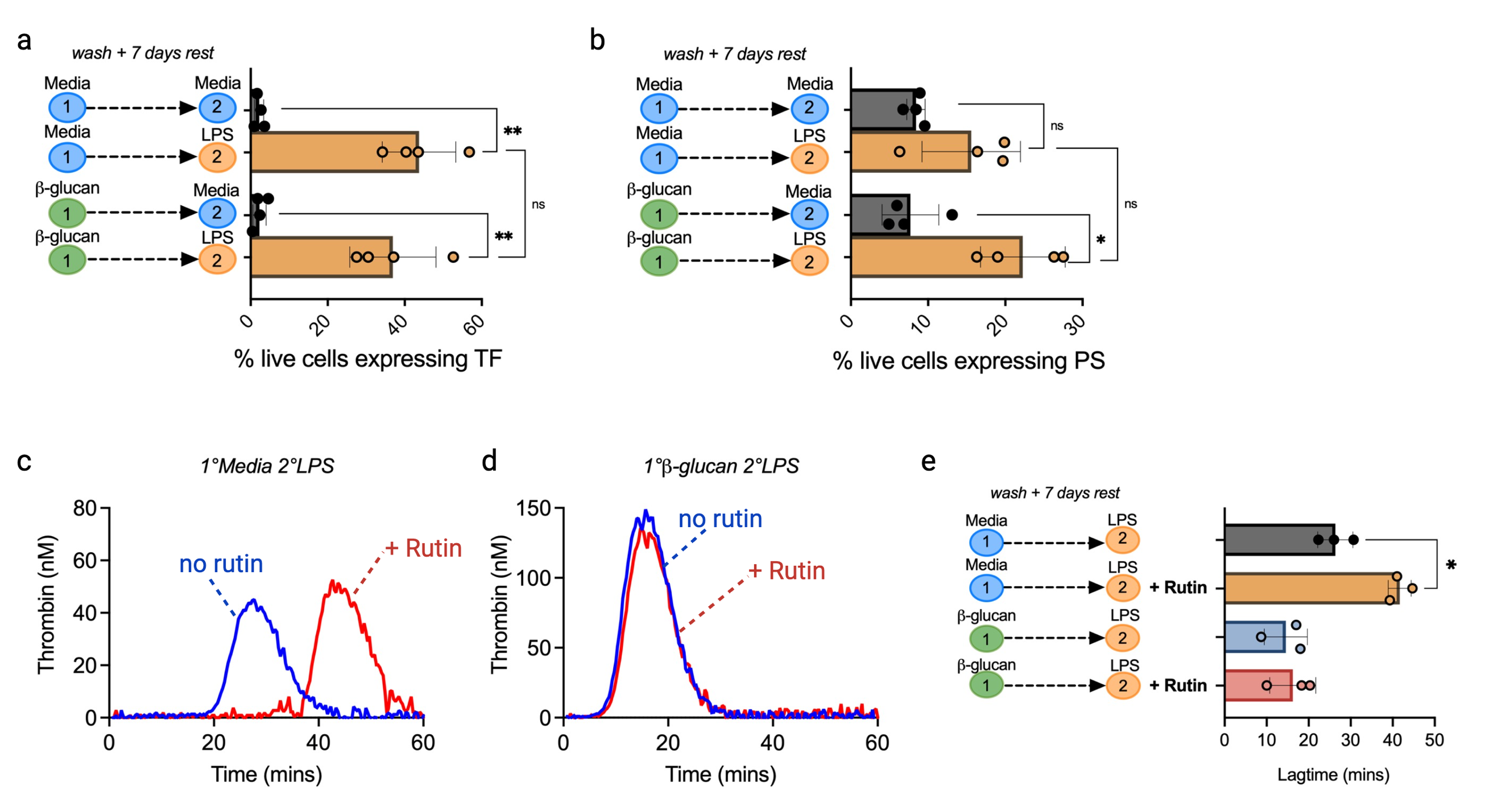

**Supplementary Figure 7**

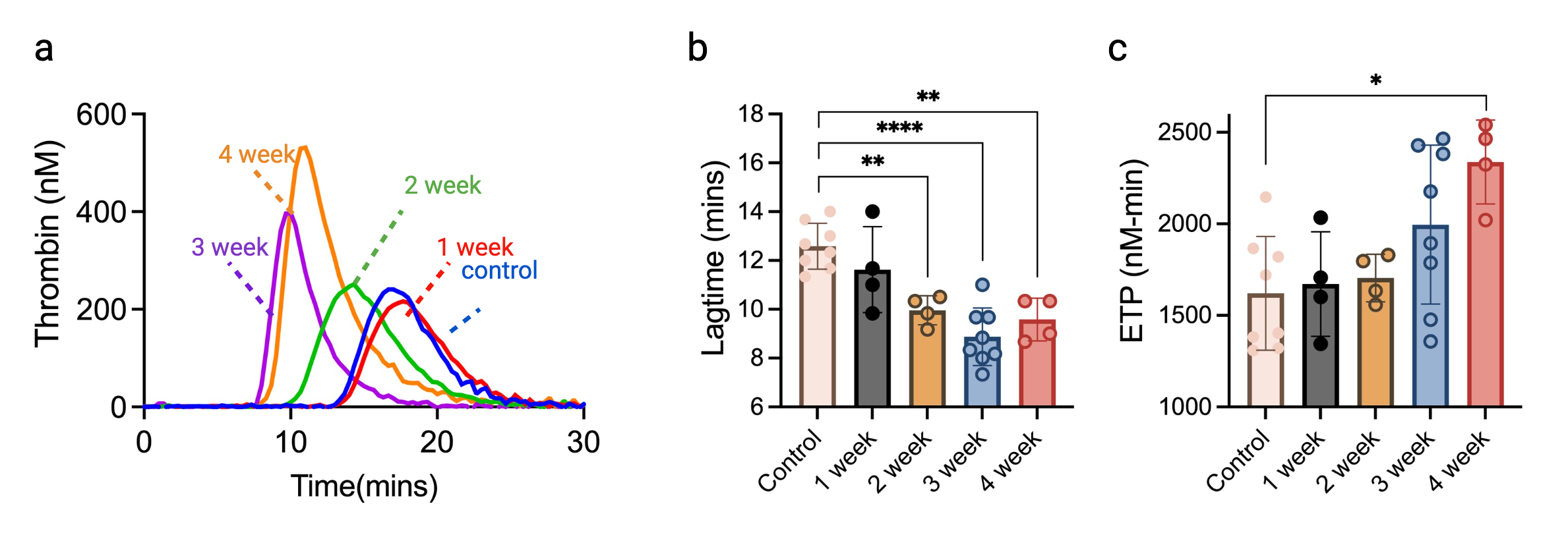

**Supplementary Figure 8**

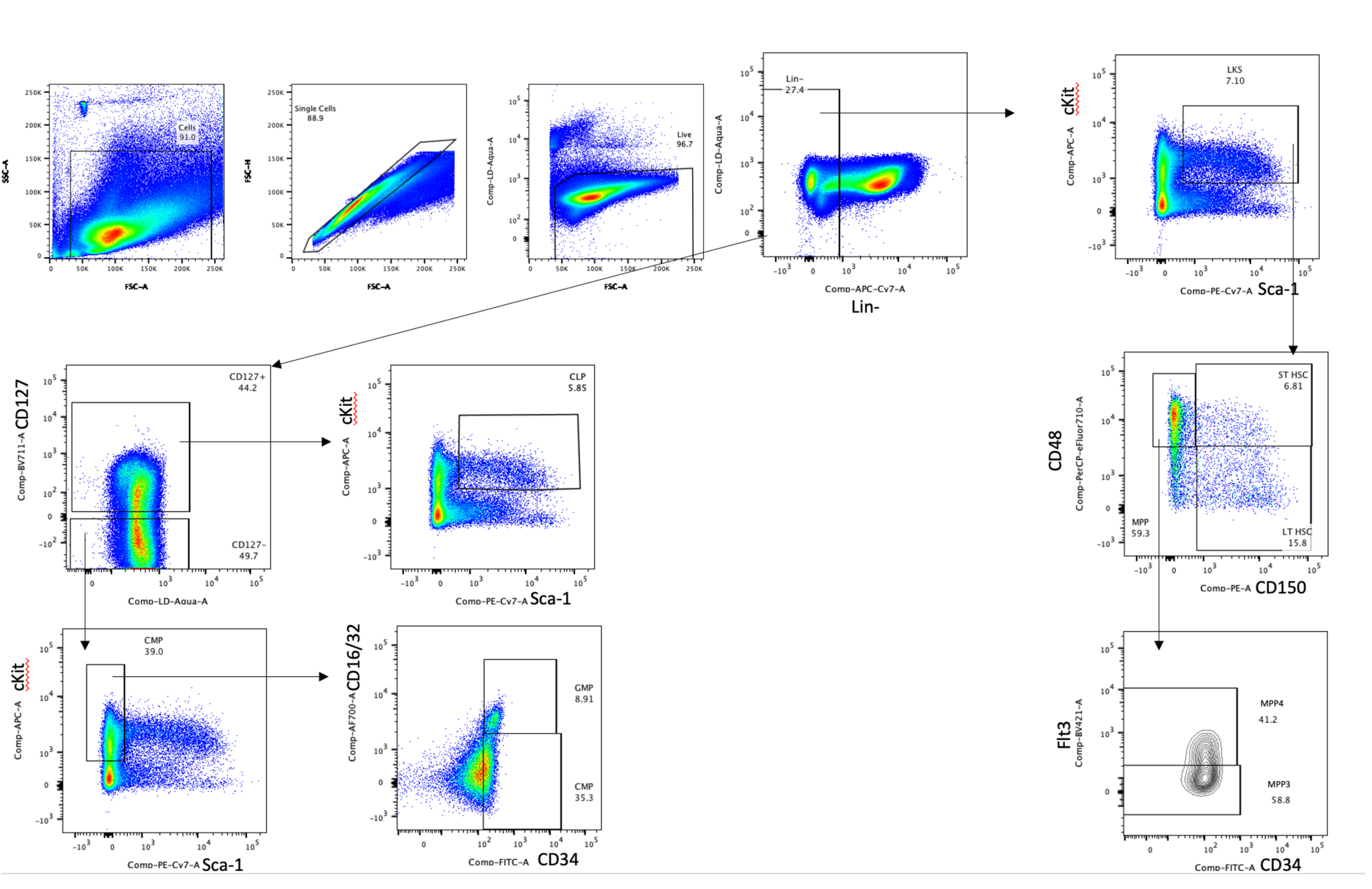

**Supplementary Figure 9**

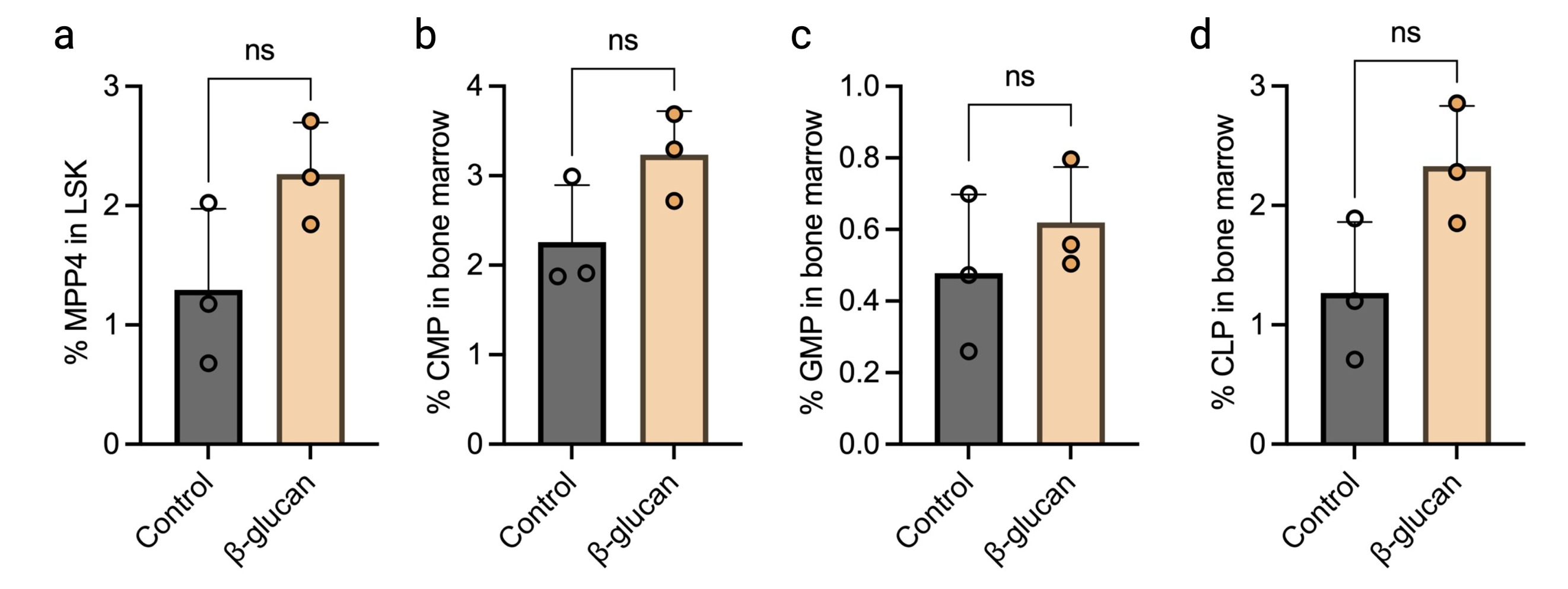

**Supplementary Figure 10**

**Supplementary Table 1: RT-qPCR primer sequences for murine and human genes**

| Primer |  | Sense |  |  | Antisense |  |
| --- | --- | --- | --- | --- | --- | --- |
| MURINE GENES |  |  |  |  |  |  |
| *RPS18* | 5’ | CTTAGAGGGACAAGTGGCG | 3’ | 5’ | ACGCTGAGCCAGTCAGTGTA | 3’ |
| *PROCR* | 5’ | CAGGACACTTGTGTGGAGTT | 3’ | 5’ | CCAGGACCAGTGATGTGTAAG | 3’ |
| *F10* | 5’ | GAAGAGCAGAACTCAGTGGTGTG | 3’ | 5’ | CAAGCTCACTGTCGCTGGTGTT | 3’ |
| *F3* | 5’ | GCACCGAGCAATGGAAGAGTTTC | 3’ | 5’ | CTTTCTGTCCCGCTCGGTTCTT | 3’ |
| *SERPINC1* | 5’ | TCCGACGCATTCCACAAA | 3’ | 5’ | GGCCAGTAATCACGACAGAA | 3’ |
| *F7* | 5’ | CAGCCCACTGCTTCGATAATA | 3’ | 5’ | GTGTCACCCGTCGTACTTG | 3’ |
| *F11* | 5’ | GATTACCAAGCACACGCATTAC | 3’ | 5’ | AAGGACGGATGGCAGAATAC | 3’ |
| *F9* | 5’ | GCATTCTGTGGAGGTGCCATCA | 3’ | 5’ | TCTCCTTTGTTCTGTGTCTTCCTT | 3’ |
| *THBD* | 5’ | GGAGAATGGTGGCTGTGAGTAC | 3’ | 5’ | GCACGATTGAACCACAGGTCTTG | 3’ |
| *TFPI* | 5’ | AGGGAACGAGAACCGATTTG | 3’ | 5’ | TGCCTTCACAGCTGTCTTC | 3’ |
| *PROS1* | 5’ | TGGCAAGGAGACAGGTGTCAGT | 3’ | 5’ | GAGCAGTGGTAACTTCCAGGAG | 3’ |
| *F2* | 5’ | ACCTTGGGACTGTGAATGTC | 3’ | 5’ | GATGGGTGGTGGAGTTGATT | 3’ |
| *F8* | 5’ | GGCGAGTAGAATGCCTTATTGGC | 3’ | 5’ | ATCACGGATGCTTCCAGAAGCC | 3’ |
| *F5* | 5’ | TGATGCTGTCCAGCCCAATAGC | 3’ | 5’ | CGATCAAGCCTGAGTGGATGTC | 3’ |
| *SERPINE1* | 5’ | CCTCTTCCACAAGTCTGATGGC | 3’ | 5’ | GCAGTTCCACAACGTCATACTCG | 3’ |
| *EGR1* | 5’ | AGCGAACAACCCTATGAGCACC | 3’ | 5’ | ATGGGAGGCAACCGAGTCGTTT | 3’ |

**Supplementary Table 2: RNAseq Differential expression of genes (DEG) analysis results for β-glucan trained BMDMs (1°β-glucan 2°LPS) compared to LPS-treated BMDMs (1°Media 2°LPS)**

| gene_id | log2FoldChange | pvalue | padj | gene_name | status |
| --- | --- | --- | --- | --- | --- |
| ENSMUSG00000021822 | 3.1286 | 2.21e-11 | 8.98e-9 | *Plau* | Up-regulated |
| ENSMUSG00000079293 | 2.1111 | 7.04e-10 | 1.82e-7 | *Clec7a* | Up-regulated |
| ENSMUSG00000027834 | 2.4216 | 8.17e-9 | 1.73e-6 | *Serpine1* | Up-regulated |
| ENSMUSG00000022126 | 1.1916 | 2.48e-9 | 5.8e-7 | *Acod1* | Up-regulated |
| ENSMUSG00000038418 | 1.2857 | 4.32e-7 | 4.84e-5 | *Egr1* | Up-regulated |
| ENSMUSG00000024401 | 1.1995 | 5.96e-7 | 6.42e-5 | *Tnf* | Up-regulated |
| ENSMUSG00000025746 | 1.8198 | 1.69e-4 | 0.00563 | *Il6* | Up-regulated |
| ENSMUSG00000028128 | 2.9656 | 4.12e-5 | 0.00195 | *F3* | Up-regulated |
| ENSMUSG00000027611 | 1.3692 | 0.00254 | 0.04093 | *Procr* | Up-regulated |
| ENSMUSG00000031444 | 1.7185 | 0.00473 | 0.06114 | *F10* | Not-Significant |

**Supplementary Table 3: RNAseq Differential expression of genes (DEG) analysis results for LPS-treated BMDMs (1°Media 2°LPS) compared to untreated BMDMs (1°Media 2°Media)**

| **gene_id** | **log2FoldChange** | **pvalue** | **padj** | **gene_name** | **status** |
| --- | --- | --- | --- | --- | --- |
| ENSMUSG00000027611 | 4.64034442 | 8.43E-95 | 4.70E-93 | *Procr* | Up-regulated |
| ENSMUSG00000024401 | 5.4380995 | 4.03E-81 | 1.75E-79 | *Tnf* | Up-regulated |
| ENSMUSG00000031444 | 3.57005048 | 2.71E-20 | 2.34E-19 | *F10* | Up-regulated |
| ENSMUSG00000038418 | 2.5470425 | 8.30E-08 | 3.15E-07 | *Egr1* | Up-regulated |
| ENSMUSG00000027834 | 0.93246693 | 0.00838 | 0.01643 | *Serpine1* | Up-regulated |
| ENSMUSG00000028128 | 1.05307692 | 0.02081 | 0.03768 | *F3* | Up-regulated |
| ENSMUSG00000025746 | -0.9012695 | 0.00349 | 0.00733 | *Il6* | Down-regulated |
| ENSMUSG00000079293 | -1.9342021 | 4.05E-06 | 1.28E-05 | *Clec7a* | Down-regulated |
| ENSMUSG00000021822 | -3.3089008 | 3.55E-12 | 1.91E-11 | *Plau* | Down-regulated |
